# Protective Effect of 2-Hydroxyestrone and 2-Hydroxyestradiol Against Chemically-Induced Hepatotoxicity *In Vitro* and *In Vivo*

**DOI:** 10.1101/2024.05.26.595989

**Authors:** Xi Sun, Xiangyu Hao, Yichen Jia, Qi Zhang, Yanyin Zhu, Yong Xiao Yang, Bao Ting Zhu

## Abstract

Ferroptosis is a form of regulated cell death closely associated with glutathione depletion and accumulation of reactive lipid peroxides. In the present study, we seek to determine whether 2-hydroxyestrone (2-OH-E_1_) and 2-hydroxyestradiol (2-OH-E_2_), two major metabolites of endogenous estrone and 17β-estradiol formed in liver by cytochrome P450 enzymes, can strongly protect against erastin– and RSL3-induced ferroptosis in hepatoma cells (H-4-II-E and HuH-7) *in vitro* and acetaminophen-induced mouse liver injury *in vivo*. We find that 2-OH-E_1_ and 2-OH-E_2_ can protect, in a dose-dependent manner, H-4-II-E hepatoma cells against erastin/RSL3-induced ferroptosis. Similar protective effect of 2-OH-E_1_ and 2-OH-E_2_ against RSL3-induced ferroptosis is also observed in HuH-7 hepatoma cells. These two estrogen metabolites strongly reduce the levels of erastin– and/or RSL3-induced accumulation of cellular NO, ROS and lipid-ROS. Mechanistically, 2-OH-E_1_ and 2-OH-E_2_ protect cells against chemically-induced ferroptosis mainly by binding to cellular protein disulfide isomerase (PDI), and then inhibit its catalytic activity and reduce PDI-catalyzed formation of iNOS dimer, thereby abrogating cellular NO, ROS and lipid-ROS accumulation. Animal studies show that 2-OH-E_1_ and 2-OH-E_2_ can also exert a strong protection against acetaminophen-induced liver injury in mice. Interestingly, while estrone and 17β-estradiol display a very weak protective effect in cultured hepatoma cells, they exert a similarly-strong protective effect as 2-OH-E_1_ and 2-OH-E_2_ *in vivo*, suggesting that the metabolic conversion of estrone and 17β-estradiol to 2-OH-E_1_ and 2-OH-E_2_ contributes importantly to their hepatoprotective effect. The results of this study reveal that 2-OH-E_1_ and 2-OH-E_2_ are important endogenous factors for protection against chemically-induced liver injury *in vivo*.

## INTRODUCTION

Ferroptosis is a form of iron-dependent regulated cell death (1,2) and is morphologically distinct from apoptosis-associated changes (1–3). Biochemically, it is usually associated with glutathione (GSH) depletion and/or inhibition of glutathione peroxidase 4 (GPX4) activity, which then leads to accumulation of lipid reactive oxygen species (ROS) and ultimately, iron-dependent oxidative cell death (ferroptosis) (1,2). Erastin, a selective inhibitor of the cystine-glutamate antiporter (system Xc^⍰^), induces ferroptosis by blocking the influx of extracellular cystine. Depletion of cellular cystine leads to blockage of intracellular synthesis of cysteine and GSH, ultimately resulting in GSH depletion (4,5). RSL3, another commonly-used prototypical inducer of ferroptosis, can inhibit GPX4 function which then leads to accumulation of lipid-ROS and ultimately ferroptotic cell death (4,6). Many studies have shown that ferroptosis is an important form of regulated cell death that is closely associated with many human diseases, such as ischemic heart disease, ischemic brain injury and renal failure, chemically-induced hepatotoxicity, and many other forms of human diseases (7–9). In addition, there is evidence suggesting that ferroptosis is linked to oxidative neurodegeneration (8,9).

Protein disulfide isomerase (PDI or PDIA1) is the prototype of the PDI family proteins, which is a ubiquitous dithiol/disulfide oxidoreductase of the thioredoxin superfamily (10,11). PDI is primarily localized in the endoplasmic reticulum of mammalian cells, although a small fraction of this protein is also found in the nucleus, mitochondria, cytosol, plasma membrane and extracellular space (11). PDI is involved in protein processing by catalyzing the formation of intra– and inter-molecular disulfide bridges in proteins (11). PDI in its reduced form can be readily converted to its oxidized form (*i.e.*, with a disulfide bond formed in the active site), which is catalyzed by the endoplasmic reticulum oxidoreductin 1 (ERO1) (12), and the oxidized PDI can be reduced by thioredoxin reductase 1 (TrxR1) (13).

In our recent studies, we have shown that PDI is involved in mediating chemically-induced ferroptosis in immortalized HT22 mouse hippocampal neuronal cells by catalyzing NOS dimerization, which is coupled with cellular nitric oxide (NO) accumulation occurring first, then followed by accumulation of cellular ROS and lipid-ROS to lethal levels, and ultimately ferroptotic cell death (14). In addition, earlier we have shown that PDI is required for catalyzing iNOS dimerization during glutamate-induced, GSH depletion-associated oxidative cytotoxicity in HT22 cells (15). Inhibition of PDI effectively abrogated glutamate-induced NOS dimerization, which was associated with a strong protection against oxidative cell death (14,15). These lines of evidence highlight that PDI is an important mediator of chemically-induced, GSH depletion-associated ferroptotic cell death, and also a potential target for cytoprotection against ferroptosis (14–16).

17β-Estradiol (E_2_) and estrone (E_1_) (structures shown in **Fig. 1**) are two most common forms of endogenous estrogens in humans. Estrogens are eliminated from the body by metabolic conversion to hormonally-inactive (or less active) water-soluble metabolites that are excreted in the urine and/or bile. Specifically, the metabolic disposition of estrogens includes oxidative metabolism and conjugative metabolism (such as glucuronidation, sulfonation and/or *O*-methylation (17,18). Members of the cytochrome P450 family are the major enzymes catalyzing NADPH-dependent oxidative metabolism of endogenous estrogens (E_2_ and E_1_) to various hydroxylated or keto metabolites in humans (19,20) and rodents (18). In the livers of humans and rodents, 2-hydroxylation of E_1_ and E_2_ to 2-OH-E_1_ and 2-OH-E_2_ (structures shown in **Fig. 1**), respectively, represents a major metabolic pathway, whereas 4-hydroxylation to 4-OH-E_2_ and 4-OH-E_1_ represents a quantitatively-minor pathway (usually <15% of 2-hydroxylation) in this organ (18–20).

**Figure 1.**
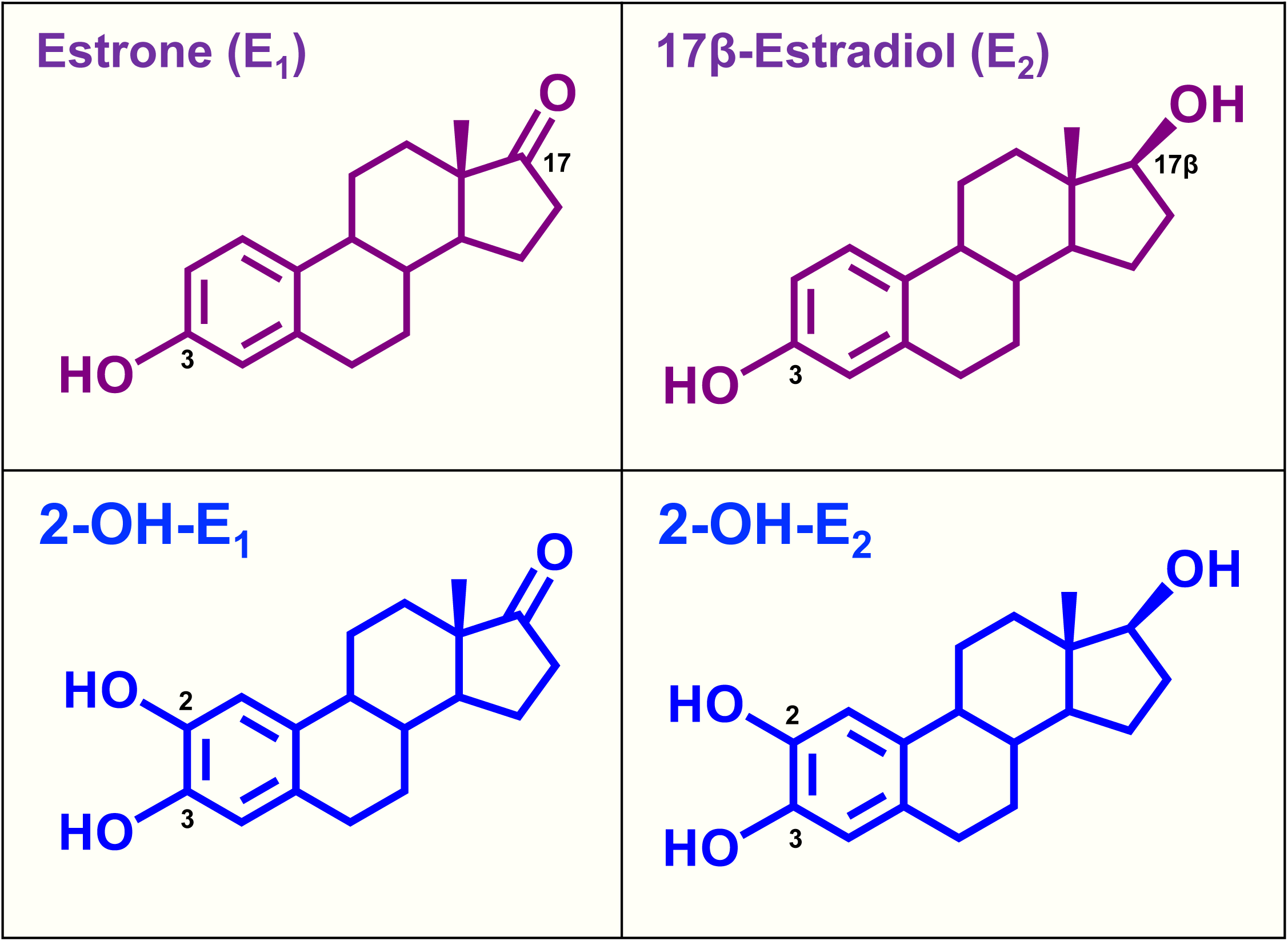
Chemical structures of estrone (E_1_), 17β-estradiol (E_2_) and their 2-hydroxylation metabolites: 2-hydroxyestrone (2-OH-E_1_) and 2-hydroxyestradiol (2-OH-E_2_).

Earlier studies have shown that some of the endogenously-formed estrogen metabolites exhibit certain biological functions that are not mediated by the classical estrogen receptor α and β (18,21,22). For instance, earlier we have shown that some of the estrogen metabolites have a strong neuroprotective effect *in vitro* and *in vivo* (21), which is independent of the estrogen receptors. In the present study, we sought to determine whether 2-OH-E_1_ and 2-OH-E_2_, two major estrogen metabolites formed in the liver, can effectively protect against erastin– or RSL3-induced ferroptosis in two hepatoma cell lines (*i.e.*, rat H-4-II-E and human HuH-7) *in vitro* and acetaminophen (APAP)-induced mouse liver injury *in vivo*. We find that 2-OH-E_1_ and 2-OH-E_2_ have a strong protective effect against chemically-induced ferroptosis both *in vitro* and *in vivo*. Mechanistically, 2-OH-E_1_ and 2-OH-E_2_ can selectively target and inhibit PDI’s catalytic activity, which is associated with abrogation of chemically-induced NOS dimerization and accumulation of cellular NO, ROS and lipid-ROS, and ultimately, a strong protection against ferroptotic cell death. The results of this study provide a novel, estrogen receptor-independent mechanism of cytoprotection against ferroptosis by two quantitatively-major endogenous estrogen metabolites.

## MATERIALS AND METHODS

### Chemicals and reagents

E_1_ and E_2_ were obtained from Sigma-Aldrich (St. Louis, MO, USA). 1,3,5(10)-Estratrien-2,3-diol-17-one (2-hydroxyestrone or 2-OH-E_1_) and 1,3,5(10)-estratrien-2,3,17β-triol (2-hydroxyestradiol or 2-OH-E_2_) were obtained from Cayman Chemical (Ann Arbor, MI, USA). E_1_, E_2_, 2-OH-E_1_ and 2-OH-E_2_ were dissolved in pure ethanol to prepare their stock solutions (usually at 10 or 20 mM). Erastin and RSL3 were purchased from Selleck Chemicals (Houston, TX, USA) and dissolved in dimethyl sulfoxide (DMSO) to prepare their stock solutions.

2’,7’-Dichlorodihydrofluorescein diacetate (DCFH-DA) and 3-amino-4-aminomethyl-2’,7’-difluorescein (DAF-FM-DA) were purchased from Beyotime Biotechnology (Shanghai, China), and BODIPY-581/591-C11 from ThermoFisher (Waltham, MA, USA). The Calcein-AM/PI Cell Viability/Cytotoxicity Assay Kit (#C2015M) was purchased from Beyotime Biotechnology (Shanghai, China), the Glutamic Pyruvic Transaminase (GPT) Kit (#C009-2-1) and the Glutamic Oxalacetic Transaminase (GOT) Kit (#C010-2-1) from Nanjing Jiancheng Bioengineering Institute (Nanjing, China), and the Malondialdehyde (MDA) Content Assay Kit (#BC0025) from Solarbio (Beijing, China). Most of the other chemicals used in this study, unless specified otherwise, were obtained from Sigma-Aldrich (Saint Louis, Missouri, USA).

### Cell culture and cell viability assay

The H-4-II-E rat hepatoma cells were obtained from the Cell Bank of the Chinese Academy of Sciences (Shanghai, China), and the human HuH-7 hepatoma cells were obtained from Otwo Biotech (Shenzhen, China). The cells were maintained in DMEM medium (for both H-4-II-E and HuH-7 cells) supplemented with 10% (*v*/*v*) fetal bovine serum (FBS; ThermoFisher, Waltham, MA, USA) and antibiotics (containing 10011U/mL penicillin and 100 μg/mL streptomycin; Sigma-Aldrich). Cells were cultured at 37LJ under 5% CO_2_, and were used for experiments when they reached 501180% confluence (depending on the types of experiments) and were usually under 25 passages. Authentication of the cell lines used in this study was performed using STR profiling and routine mycoplasma testing.

For cell viability assays, cells were seeded in 96-well plates at a density of 2,000 cells/well for H-4-II-E cells and 5000 cells/well for HuH-7 cells, and treated with different drugs as indicated. 3-(4,5-Dimethylthiazol-2-yl)-2,5-diphenyltetrazolium bromide (MTT, MedChemExpress, Monmouth Junction, NJ, USA) was added to each well at a final concentration of 0.5 mg/mL and incubated for 3 h at 37°C under 5% CO_2_. After incubation, DMSO was added to dissolve the MTT formazan, and absorbance was measured at 560 nm wavelength with a microplate reader (Biotek, Winooski, VT, USA).

### Measurement of cellular NO and ROS levels by fluorescence microscopy

The cells were plated in 24-well plates at a density of 5 lZ 10^4^ per well, and then given different drug treatments as indicated. For staining of cellular NO and total ROS, the cells were first washed twice with HBSS and then incubated with DAF-FM-DA (5 μM, for staining cellular NO) and DCFH-DA (5 μM, for staining of cellular ROS), respectively, in 200 μL DMEM (serum-free and phenol red-free) for 20 min at 37°C. Following three washes with HBSS, fluorescence images were captured using an AXIO fluorescence microscope (Carl Zeiss Corporation, Germany).

### Confocal microscopy

For visualization of subcellular distribution of cellular lipid-ROS, the cells were seeded at a density of 5 lZ 10^4^ per well on coverslips placed inside the 24-well plates. Twenty-four h later, cells were treated with drugs as indicated. Coverslips were then washed in HBSS and incubated in HBSS containing BODIPY-581/591-C11 (5 μM) for 20 min at 37°C under 5% CO_2_. Coverslips were then mounted on microscope slides for visualization. Slides were imaged using a LSM-900 confocal laser scanning microscope (Carl Zeiss), and images were analyzed with the Zen software (Carl Zeiss).

### Flow cytometry

For quantitative analysis of cellular levels of NO, ROS and lipid-ROS, the cells were plated in 6-well plates at a density of 15 × 10^4^ cells/well for 24 h before the start of drug treatments. Following treatment with different drugs as indicated, cells were trypsinized, collected and suspended in phosphate-buffered saline (PBS). Cells were then centrifuged, and the resulting cell pellets were resuspended in DMEM (free of phenol red and serum) containing DAF-FM-DA (5 μM, for staining of cellular NO), DCFH-DA (5 μM, for staining of cellular total ROS) and BODIPY-581/591-C11 (5 μM, for staining of cellular lipid-ROS), respectively. After a 20-min incubation at 37°C, the cells were washed three times with HBSS to remove any remaining fluorescent dyes. Levels of cellular NO, ROS and lipid-ROS were measured by a flow cytometer (Beckman Coulter, Brea, CA, USA) and analyzed using the FlowJo software (FlowJo, LLC, Ashland, USA).

### Western blot analysis

Western blot analysis was performed as described previously (58) using the following primary antibodies: rabbit anti-iNOS (#ab178945; Abcam, Cambridge, MA, USA), mouse anti-β-actin (#GB113225; Servicebio, Wuhan, China) and rabbit anti-PDI (#3501S; Cell Signaling Technology, Beverly, MA, USA). The mouse anti-rabbit (#3678S) and the anti-mouse HRP-conjugated secondary antibodies (#7076S) were from Cell Signaling Technology (Beverly, MA, USA). Enhanced chemiluminescence was used for detection (Amershan Bioscience, Freiburg, Germany). The experiments were repeated multiple times to confirm the observations. Representative blots from one representative experiments are shown.

### Cellular thermal shift assay (CETSA)

CETSA was conducted according to the protocol described earlier (59,60) to determine whether 2-OH-E_1_ and 2-OH-E_2_ can bind to the PDI protein in live cells in culture. Briefly, after the cells were treated with 2-OH-E_1_ or 2-OH-E_2_ or vehicle for 3 h, they were washed with ice-cold PBS, harvested by trypsinization, centrifuged, and resuspended in PBS supplemented with the complete protease inhibitor cocktail (Selleck Chemicals, Houston, TX, USA). Equal amounts of cell suspensions were aliquoted into 0.2 mL PCR microtubes and incubated for 30 min at room temperature. Subsequently, aliquots of the cell suspension were heated individually in a Ristretto Thermal Cycler (VWR, Darmstadt, Germany) at the indicated temperatures for 5 min, followed by cooling for 5 min on ice. Finally, the cells were lysed using three cycles of freeze–thawing, and the soluble fractions were isolated by centrifugation and analyzed by SDS-PAGE followed by Western blotting as described above.

### Assay of PDI catalytic activity and its inhibition by catechol estrogens

The effect of 2-OH-E_1_ and 2-OH-E_2_ on the catalytic activity of purified wild-type PDI and mutant PDI-Ala256 was determined by analyzing PDI-catalyzed aggregation of the insulin B chain as described earlier with some modifications (61,62). Briefly, insulin (125 μM) was incubated in a 96-well plate in 10 mM sodium phosphate buffer (pH 7.4) and 5 mM DTT with or without the recombinant wild-type PDI or mutant PDI-Ala256 (0.2 μg/μL). The aggregation was monitored at 37°C using a Synergy Plate Reader (Biotek, Winooski, VT, USA), with the wavelength set at 650 nm.

### Molecular docking analysis

The PDI-ligand interactions were analyzed using the molecular docking method. The structures of human PDI (PDB code **6I7S**; chain A) (23) and E_1_ (PDB code **6MNE**; ligand ID **J3Z**) (24) and E_2_ (PDB code **6KEP**; ligand ID **EST**) (25) were downloaded from the Protein Data Bank (https://www.rcsb.org/) (26). The structures of 2-OH-E_1_ and 2-OH-E_2_ were generated based on the structures of E_1_ and E_2_ by adding the hydroxyl group. The experimental structures of PDI and the generated structures of 2-OH-E_1_ and 2-OH-E_2_ were adopted as the receptor and ligands, respectively.

The structures of PDI, 2-OH-E_1_ and 2-OH-E_2_ were preprocessed using the Protein Preparation Wizard in Schrodinger Suite (Maestro 12.8, 2021; Schrodinger LLC, New York, NY, USA). The hydrogen atoms were added, and the PDI structure was optimized using OPLS4 force field (27). The PDI–ligand docking decoys were generated using Glide-XP (extra precision) in Schrodinger Glide software (28). The nearby torsional minima of the lowest energy PDI–ligand binding poses were sampled using the Monte Carlo (MC) procedure. The Cα atom of His256 in the b’ domain of PDI was employed as the center of the docking grid box with dimensions set at 30 × 30 × 30 Å^3^.

The representative scoring function of PRODIGY-LIG (29) was used for further filtering of the docking results. PRODIGY-LIG is an empirical scoring function that was developed to calculate protein–ligand binding affinity largely based on the number of atomic contacts and electrostatic energy (29).

To investigate the importance of His256 in the PDI-ligand interactions, His256 was mutated to alanine (Ala) in the representative predicted structures of the PDI–ligand complexes. The relative binding energies or affinities were predicted using PRODIGY-LIG (29).

Molecular dynamics simulations were used to generate more representative conformations for the wild-type PDI–2-OH-E_1/2_ complexes. These representative conformations were used for further comparison of the binding affinities between the wild-type PDI and 2-OH-E_1/2_. The complexes were preprocessed to generate the topology files using *CHARMM-GUI* (https://charmm-gui.org/) (30). The force field parameters of 2-OH-E_1/2_ were generated based on *CHARMM* general force field (31), and those of the wild-type PDI protein based on *CHARMM36m* force field (32). The systems were embedded into a rectangular water box which extends the solvent 10 Å in x, y, z directions, and the TIP3P water model (33) was adopted. K^+^ and Cl^−^ ions with parameters approximated by Roux et al. (34) were employed to neutralize the systems’ charges. The energy minimization (10000 steps), equilibrium simulation (0.25 ns) in NVT ensemble and production simulation (100 ns) in NPT ensemble were conducted using NAMD (35). The time step was set to 2 fs, and the temperature was maintained at 300 K using Langevin dynamics (36), respectively. The periodic boundary conditions were adopted, the short-range electrostatic and van der Waals interactions were truncated smoothly with a cutoff (12 Å) and a switching function was adopted at 10 Å. Long-range electrostatic interaction was estimated by the particle mesh Ewald algorithm (37,38). The pressure in NPT ensemble was maintained at 1 atm by the Langevin piston method (39). The binding affinities/energies in the representative conformations were predicted using PRODIGY-LIG (29).

### Animal experiments

All procedures involving the use of live animals described in this study were approved by the Institutional Animal Care and Use Committee (IACUC) of The Chinese University of Hong Kong (Shenzhen) (October 28, 2021; CUHKSZ-AE2021002), and the guidelines for the humane care of animals set forth by the U.S. National Institutes of Health were followed.

The C57BL/6J male mice (91111 weeks old) were purchased from Charles River (China). For APAP-induced hepatocyte injury, the mice were randomly divided into different experimental groups (with 6 animals per group) with comparable average body weights. APAP was given to mice by gastric intubation at a dose of 300 mg/kg, and 2-OH-E_1_ or 2-OH-E_2_ (at the doses of 10 and 100 μg/mouse) was injected intraperitoneally (*i.p.*). The plasma levels of glutamic-pyruvic transaminase (GPT) and glutamic-oxalacetic transaminase (GOT) were determined using the GPT and GOT kits according to manufacturers’ instructions.

For histological staining of tissue slides, fresh mouse liver tissues were placed in 10% formalin solution and stored for at least three days before dehydration. The tissues were sectioned to 4-μm thickness after completion of dehydration and paraffin embedding for H/E staining, and the images were captured at 200lZ magnification with an upright microscope.

### Statistical analysis

The quantitative experiments conducted in this study were repeated multiple times to validate the experimental observations. All statistical analyses were performed with GraphPad Prism 10 software (GraphPad Software, La Jolla, CA) using one-way ANOVA followed by Dunnett’s post-hoc tests for multiple comparisons. The data were presented as the mean ± standard deviation (S.D.), typically obtained from a representative experiment with multiple replicates. Statistical significance was denoted by *P* < 0.05 (* or ^#^) and *P* < 0.01 (** or ^##^) for significant and very significant differences, respectively. In most cases, * and ** denote the comparison for statistical significance between the control group (cells treated with vehicle only) and the cells treated with a ferroptosis inducer (erastin or RSL3), whereas ^#^ and ^##^ denote the comparison between the cells treated with erastin or RSL3 and the cells jointly treated with erastin/RSL3 and another compound. For Western blot quantification, one representative data is shown.

## RESULTS

### 2-OH-E_1_ and 2-OH-E_2_ protect against chemically-induced ferroptotic cell death

In this study, the protective effect of 2-OH-E_1_ and 2-OH-E_2_ against chemically-induced ferroptosis was analyzed first in cultured cells. Specifically, two hepatoma cell lines (*i.e.*, H-4-II-E rat hepatoma cells and HuH-7 human hepatoma cells) and two commonly-used ferroptosis inducers (*i.e.*, erastin and RSL3) were jointly used to assess the cytoprotective actions of 2-OH-E_1_ and 2-OH-E_2_ *in vitro*.

### H-4-II-E cells

First, the H-4-II-E rat hepatoma cells were treated with increasing concentrations of erastin for 24 h, and the cell viability was assessed based on MTT assay and change in gross morphology. Erastin decreased the cell viability (MTT assay) in a dose-dependent manner, with a maximal reduction by ∼85% when erastin was present at 2.4–3 μM concentrations (**Fig. 2A**). Change in cellular gross morphology confirmed erastin-induced cell death (data now shown). Erastin-induced cell death could be effectively rescued by Fer-1 and Trolox, but not by z-VAD-FAM (**Fig. 2B**), which is consistent with the induction of ferroptotic cell death. It is of note that the sensitivity of H-4-II-E cells to erastin-induced cytotoxicity often varied slightly from experiments to experiments. In the subsequent experiments that were designed to determine the protective effect of 2-OH-E_1_ and 2-OH-E_2_ against erastin-induced cytotoxicity, a 1-μM concentration of erastin was used which usually induced 70–80% reduction in cell viability when present alone.

**Figure 2.**
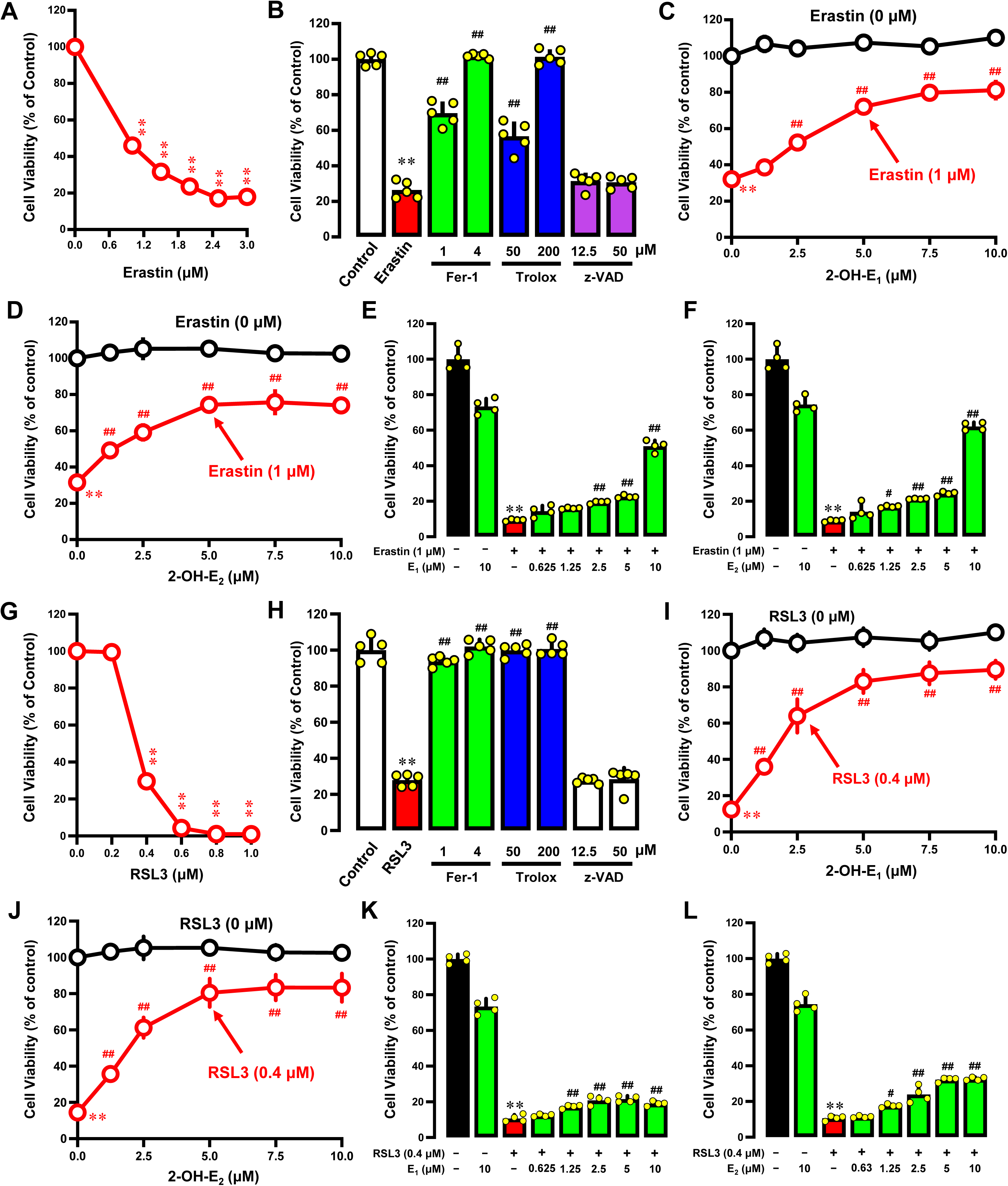
2-OH-E_1_ and 2-OH-E_2_ protect against chemically-induced ferroptotic cell death in H-4-II-E rat hepatoma cells. **A.** Dose-dependent effect of erastin on cell viability after 24-h treatment (MTT assay, n = 5). **B.** Effect of Fer-1, Trolox and z-VAD-FAM on erastin-induced cell death. Cells were treated with erastin (1 μM) ± Fer-1, Trolox or z-VAD-FAM at the indicated concentrations for 24 h, and cell viability was determined by MTT assay (n = 5). **C, D.** Protection against erastin-induced cytotoxicity by 2-OH-E_1_ (**C**) and 2-OH-E_2_ (**D**). Cells were treated with erastin (1 μM) ± 2-OH-E_1_ or 2-OH-E_2_ at the indicated concentrations for 24 h, and cell viability was determined by MTT assay (n = 5). **E, F.** Effect of E_1_ (**E**) and E_2_ (**F**) on erastin-induced cytotoxicity (MTT assay; n = 4). **G.** Dose-dependent effect of RSL3 on cell viability after 24-h treatment (MTT assay, n = 5). **H.** Effect of Fer-1, Tolox and z-VAD-FAM on RSL3-induced cell death. Cells were treated with RSL3 (0.4 μM) ± Fer-1, Trolox or z-VAD-FAM at the indicated concentrations for 24 h, and cell viability was determined by MTT assay (n = 5). **I, J.** Protection against RSL3-induced cytotoxicity by 2-OH-E_1_ (**I**) and 2-OH-E_2_ (**J**). Cells were treated with RSL3 (0.4 μM) ± 2-OH-E_1_ or 2-OH-E_2_ at the indicated concentrations for 24 h, and cell viability was determined by MTT assay (n = 5). **K, L.** Effect of E_1_ (**K**) and E_2_ (**L**) on RSL3-induced cytotoxicity (MTT assay, n = 4). All quantitative data are presented as mean ± S.D. (* or ^#^ *P* < 0.05; ** or ^##^ *P* < 0.01).

As shown in **Fig. 2C**, the presence of 2-OH-E_1_ alone at 1.25–12.5 μM was not cytotoxic in H-4-II-E cells. Joint treatment of cells with erastin + 2-OH-E_1_ exerted a dose-dependent protection against erastin-induced ferroptosis (**Fig. 2C**), with a maximal protection of ∼80% when 2-OH-E_1_ was present at 7.5–10 μM. Similarly, the presence of 2-OH-E_2_ alone at 1.25–10 μM was not cytotoxic, but joint treatment of cells with erastin + 2-OH-E_2_ exerted a similar dose-dependent protection against erastin-induced ferroptosis (**Fig. 2D**). The maximal protection (74%) was seen when 2-OH-E_2_ was present at 5–7.5 μM. It is of note that E_1_ and E_2_ at the same concentrations had a markedly weaker protective effect against erastin-induced ferroptosis in H-4-II-E cells (**Fig. 2E, 2F**).

In this study, we also determined the protective effect of 2-OH-E_1_ and 2-OH-E_2_ against RSL3-induced ferroptotic cell death in H-4-II-E cells. Treatment of cells with RSL3 alone caused a dose-dependent reduction in cell viability (**Fig. 2G**). RSL3-induced cell death in H-4-II-E cells was effectively rescued by Fer-1 and Trolox, but not by z-VAD-FAM (**Fig. 2H**), confirming the induction of ferroptotic cell death. At 0.4 µM, RSL3 reduced the cell viability by ∼70%, and this concentration was used in subsequent experiments to determine the cytoprotective effect of 2-OH-E_1_ and 2-OH-E_2_.

Joint treatment of cells with RSL3 + 2-OH-E_1_ exerted a dose-dependent protection against RSL3-induced ferroptosis, and a maximal protection of 88% was observed when 2-OH-E_1_ was present at 7.5–10 μM (**Fig. 2I**). Similarly, joint treatment of cells with RSL3 +2-OH-E_2_ also exerted a dose-dependent protection against RSL3-induced ferroptosis, and the maximal protection (83%) was observed when 2-OH-E_2_ was present at 5–10 μM (**Fig. 2J**). Notably, E_1_ and E_2_ only displayed a very weak protective effect against RSL3-induced ferroptosis in H-4-II-E cells (**Fig. 2K, 2L**).

### HuH-7 cells

The human HuH-7 hepatoma cells were highly sensitive to RSL3-induced ferroptosis. Based on MTT assay and analysis of cellular gross morphology, RSL3 reduced the viability of HuH-7 cells in a concentration-dependent manner (**Fig. 3A**). RSL3-induced cell death could be effectively rescued by Fer-1 and Trolox, but not by z-VAD-FAM (**Fig. 3B**), consistent with the induction of ferroptotic cell death. Because 0.5 μM RSL3 elicited ∼80% reduction in the cell viability, this concentration of RSL3 was used to further evaluate the cytoprotective effect of 2-OH-E_1_ and 2-OH-E_2_.

**Figure 3.**
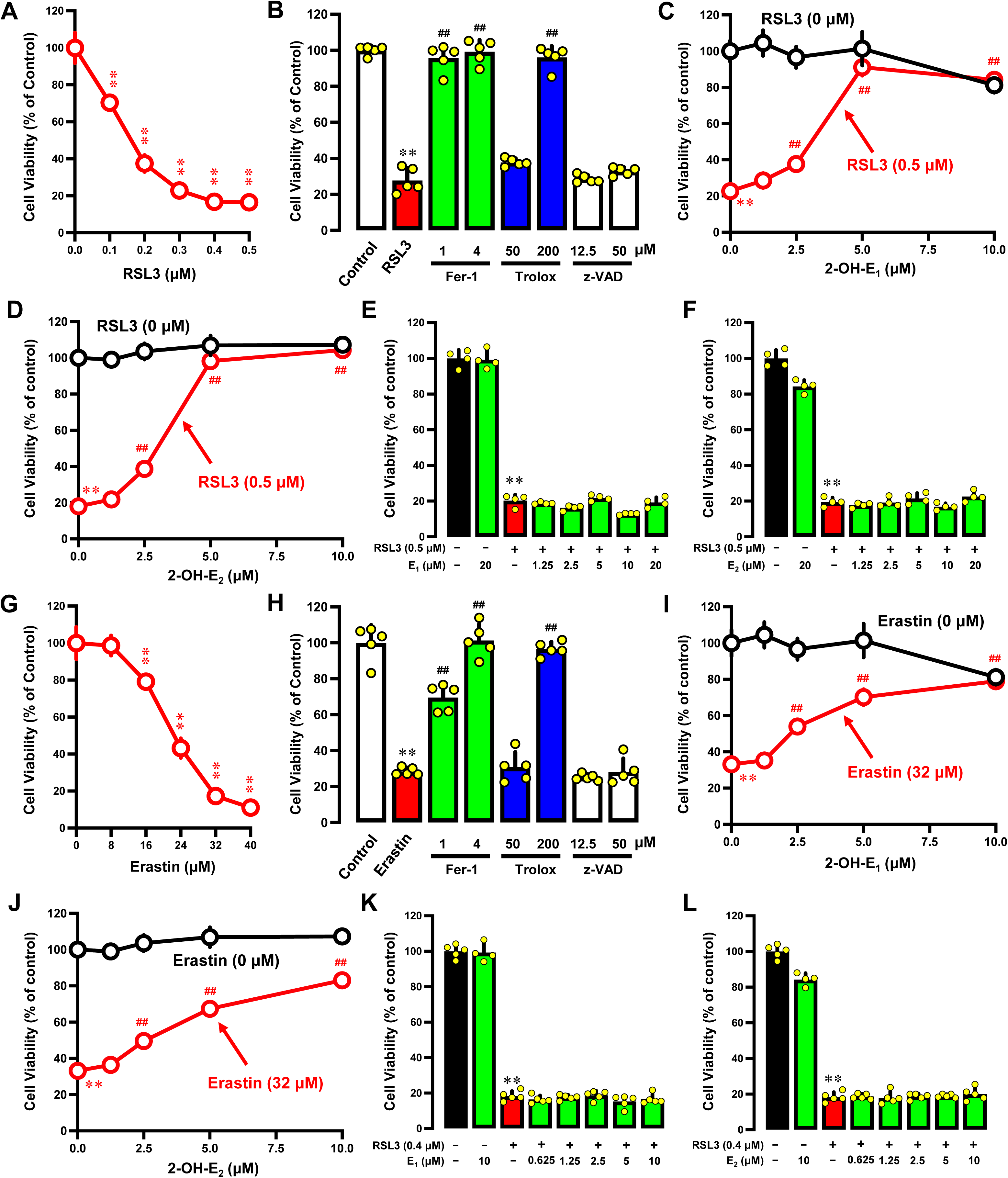
2-OH-E_1_ and 2-OH-E_2_ protect against chemically-induced ferroptotic cell death in HuH-7 human hepatoma cells. **A.** Dose-dependent effect of RSL3 on cell viability after 24-h treatment (MTT assay, n = 5). **B.** Effect of Fer-1, Trolox and z-VAD-FAM on RSL3-induced cell death. Cells were treated with RSL3 (0.5 μM) ± Fer-1, Trolox or z-VAD-FAM at the indicated concentrations for 24 h, and cell viability was determined by MTT assay (n = 5). **C, D.** Protection against RSL3-induced cytotoxicity by 2-OH-E_1_ (**C**) and 2-OH-E_2_ (**D**). Cells were treated with RSL3 (0.5 μM) ± 2-OH-E_1_ or 2-OH-E_2_ at the indicated concentrations for 24 h, and cell viability was determined by MTT assay (n = 5). **E, F.** Effect of E_1_ (**E**) and E_2_ (**F**) on RSL3-induced cytotoxicity (MTT assay; n = 4). **G.** Dose-dependent effect of erastin on cell viability after 24-h treatment (MTT assay, n = 5). **H.** Effect of Fer-1, Trolox and z-VAD-FAM on erastin-induced cell death. Cells were treated with Erastin (32 μM) ± Fer-1, Trolox or z-VAD-FAM at the indicated concentrations for 24 h, and cell viability was measured by MTT assay (n = 5). **I, J.** Protection against erastin-induced cytotoxicity by 2-OH-E_1_ (**C**) and 2-OH-E_2_ (**D**). Cells were treated with Erastin (32 μM) ± 2-OH-E_1_ or 2-OH-E_2_ at the indicated concentrations for 24 h, and cell viability was measured by MTT assay (n = 5). **K, L.** Effect of E_1_ (**E**) and E_2_ (**F**) on erastin-induced cytotoxicity (MTT assay; n = 5). All quantitative data are presented as mean ± S.D. (* or ^#^ *P* < 0.05; ** or ^##^ *P* < 0.01).

Joint treatment of cells with RSL3 + 2-OH-E_1_ exerted a concentration-dependent protection against RSL3-induced cell death, and a 90% protection was seen at 5 μM 2-OH-E_1_ (**Fig. 3C**). Similarly, 2-OH-E_2_ also exhibited a concentration-dependent strong protection against RSL3-induced ferroptosis, and the protection at 51110 μM was nearly 100% (**Fig. 3D**). Notably, treatment of cells with E_1_ or E_2_ at the same concentrations did not display a meaningful protection against 0.5 μM RSL3-induced ferroptosis (**Fig. 3E**, **3F**).

It was observed that HuH-7 human hepatoma cells were markedly less sensitive to erastin-induced cell death compared to H-4-II-E rat hepatoma cells (compare **Fig. 3G** with **Fig 2A**). However, the cell death induced by erastin (at 25 µM) in HuH-7 cells could still be effectively rescued by Fer-1 and Trolox but not by z-VAD-FMA (**Fig. 3H**). We found that 2-OH-E_1_ and 2-OH-E_2_ also elicited a protective effect against erastin-induced cell death in HuH-7 cells (**Fig. 3I, 3J**), whereas E_1_ and E_2_ had little or no protective effect (**Fig. 3K, 3L**).

Together, these results show that 2-OH-E_1_ and 2-OH-E_2_ have a strong protective effect against erastin/RSL3-induced ferroptotic cell death in both H-4-II-E and HuH-7 cells. In comparison, E_1_ and E_2_ have either a much weaker or no appreciable protective effect under the same experimental conditions.

### 2-OH-E_1_ and 2-OH-E_2_ abrogate chemically-induced accumulation of cellular NO, ROS and lipid-ROS

Our recent studies have shown, for the first time, that during the induction of ferroptosis by erastin and RSL3, the cellular NO level was increased in a time-dependent manner, and the NO buildup subsequently led to cellular ROS and lipid-ROS accumulation (14,16,40). In this study, we sought to determine whether 2-OH-E_1_ and 2-OH-E_2_ can modulate erastin– and RSL3-induced accumulation of NO, ROS and lipid-ROS in both H-4-II-E and HuH-7 cells.

### H-4-II-E cells

Treatment of cells with 1 μM erastin for 8 h increased cellular NO accumulation (**Fig. 4A**). While 2-OH-E_1_ (7.5 μM) alone did not meaningfully change cellular NO level, joint treatment of cells with erastin + 2-OH-E_1_ abrogated erastin-induced NO accumulation by 63.8% (*P* < 0.01; **Fig. 4A**). 2-OH-E_2_ had a similar protective effect against erastin-induced NO accumulation, resulting in 61.8% reduction (*P* < 0.01; **Fig. 4D**).

**Figure 4.**
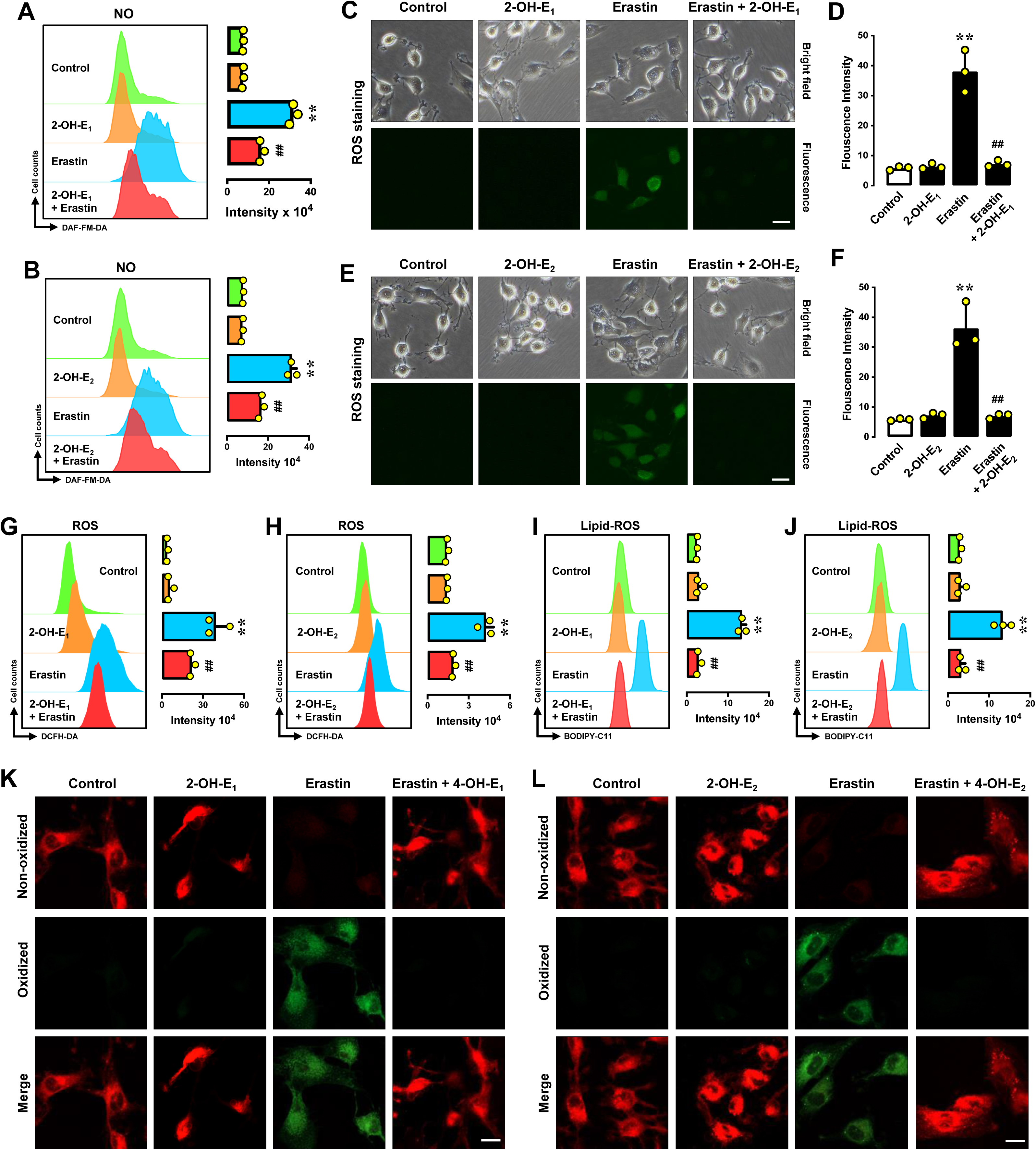
2-OH-E_1_ and 2-OH-E_2_ abrogate erastin-induced accumulation of cellular NO, ROS and lipid-ROS in H-4-II-E rat hepatoma cells. **A, B.** Cellular level of NO after 8-h treatment with erastin (1 μM) ± 2-OH-E_1_ (7.5 μM) or 2-OH-E_2_ (5 μM). The left panels of **A** and **B** are flow cytometry data, and their right panels are the respective quantitative intensity values (n = 3). **C–H.** Cellular level of ROS after 8-h treatment with 1 μM erasin ± 7.5 μM 2-OH-E_1_ or 5 μM 2-OH-E_2_. **C, D** are fluorescence microscopy images (scale bar = 60 μm), and **D**, **F** are respective quantitative intensity values (n = 3). **G, H.** The left panels are flow cytometry histograms, and the right panels are the respective quantitative intensity values (n = 3). **I–L.** Cellular levels of lipid-ROS after 8-h treatment with 1 μM erastin ± 7.5 μM 2-OH-E_1_ or 5 μM 2-OH-E_2_. The left panels of **I** and **J** are flow cytometry data, and their right panels are the respective quantitative intensity values (n = 3). **K** and **L** are confocal microscopy images (scale bar = 30 μm). All quantitative data are presented as mean ± S.D. (* or ^#^ *P* < 0.05; ** or ^##^ *P* < 0.01).

Treatment of cells with erastin alone for 8 h also increased cellular ROS accumulation (**Fig. 4B, 4C**). While 2-OH-E_1_ alone did not significantly change cellular ROS level, joint treatment of cells with erastin + 2-OH-E_1_ reduced the accumulation of cellular ROS by 49.4% (*P* < 0.05; **Fig. 4B, 4C, 4G**). A similar effect was observed with 2-OH-E_2_, abrogating erastin-induced ROS accumulation by 83.2% (*P* < 0.01; **Fig. 4E, 4F, 4H**).

Similarly, cells treated with erastin alone for 8 h displayed cellular lipid-ROS accumulation (**Fig. 4I –4L**). While treatment of cells with 2-OH-E_1_ alone did not change lipid-ROS level, joint treatment with erastin + 2-OH-E_1_ strongly reduced lipid-ROS accumulation (**Fig. 4I, 4K**). Joint treatment of cells with 2-OH-E_2_ similarly abrogated erastin-induced lipid-ROS accumulation (**Fig. 4J, 4L**).

In this study, we have also determined the effect of 2-OH-E_1_ and 2-OH-E_2_ on RSL3-induced accumulation of cellular NO, ROS and lipid-ROS. Treatment of cells with 0.4 μM RSL3 for 4 h increased the cellular NO accumulation (**Fig. 5A, 5D**), and joint treatment of cells with 2-OH-E_1_ abrogated RSL3-induced NO accumulation (**Fig. 5A**). A similar abrogation of NO accumulation was observed with 2-OH-E_2_ (**Fig. 5D**).

**Figure 5.**
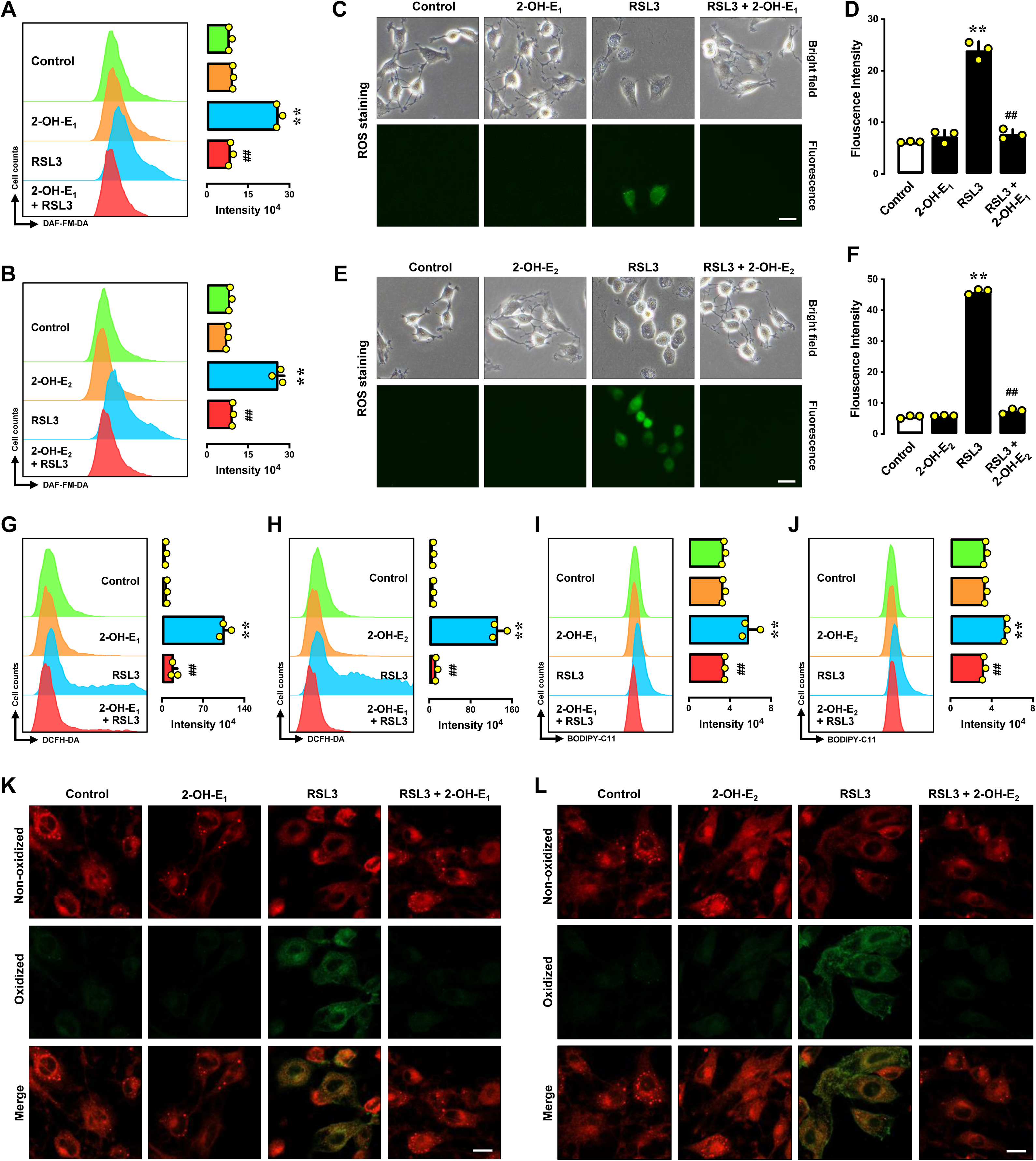
2-OH-E_1_ and 2-OH-E_2_ abrogate RSL3-induced accumulation of cellular NO, ROS and lipid-ROS in H-4-II-E rat hepatoma cells. **A, B.** Cellular level of NO after 4-h treatment with RSL3 (0.4 μM) ± 2-OH-E_1_ (7.5 μM) or 2-OH-E_2_ (5 μM). The left panels of **A** and **B** are flow cytometry data, and their right panels are the respective quantitative intensity values (n = 3). **C–H.** Cellular level of ROS after 4-h treatment with RSL3 (0.4 μM) ± 2-OH-E_1_ (7.5 μM) or 2-OH-E_2_ (5 μM). **C, D** are fluorescence microscopy images (scale bar = 60 μm), and **D**, **F** are respective quantitative intensity values (n = 3). **G, H.** The left panels are flow cytometry histograms, and the right panels are the respective quantitative intensity values (n = 3). **I–L.** Cellular level of lipid-ROS after 4-h treatment with RSL3 (0.4 μM) ± 2-OH-E_1_ (7.5 μM) or 2-OH-E_2_ (5 μM). The left panels of **I** and **J** are flow cytometry data, and their right panels are the respective quantitative intensity values (n = 3). **K** and **L** are confocal microscopy images (scale bar = 30 μm). All quantitative data are presented as mean ± S.D. (* or ^#^ *P* < 0.05; ** or ^##^ *P* < 0.01).

Treatment of cells with RSL3 alone for 4 h also significantly elevated cellular ROS accumulation (**Fig. 5B, 5C, 5E, 5F, 5G, 5H**), and joint treatment of cells with 2-OH-E_1_ abrogated RSL3-induced ROS accumulation (**Fig. 5B, 5C, 5G**). A similar pattern was observed with 2-OH-E_2_ (**Fig. 5E, 5F, 5H**). Similarly, lipid-ROS levels were also increased in cells treated with 0.4 μM RSL3 alone for 4 h (**Fig. 5I, 5J, 5K, 5L**), and joint treatment of cells with 2-OH-E_1_ strongly abrogated RSL3-induced lipid-ROS accumulation by (**Fig. 5I, 5K**). A similar trend was observed with 2-OH-E_2_ (**Fig. 5J, 5L**).

### HuH-7 cells

While treatment of HuH-7 cells with 10 μM 2-OH-E_1_ or 2-OH-E_2_ alone did not increase cellular NO levels, joint treatment of cells with RSL3 + 2-OH-E_1_ or 2-OH-E_2_ abrogated RSL3-induced NO accumulation (**Fig. 6A, 6B**). Similarly, treatment with 2-OH-E_1_ or 2-OH-E_2_ alone did not increase cellular ROS and lipid-ROS levels in HuH-7 cells, but joint treatment of cells with RSL3 + 2-OH-E_1_ or 2-OH-E_2_ abrogated RSL3-induced accumulation of cellular ROS (**Fig. 6B, 6C, 6G** for 2-OH-E_1_; **Fig. 6E, 6F, 6H** for 2-OH-E_2_) and lipid-ROS (**Fig. 6I** for 2-OH-E_1_; **Fig. 6J** for 2-OH-E_2_). The change in lipid-ROS levels was also confirmed by confocal microscopy (**Fig. 6K, 6L**).

**Figure 6.**
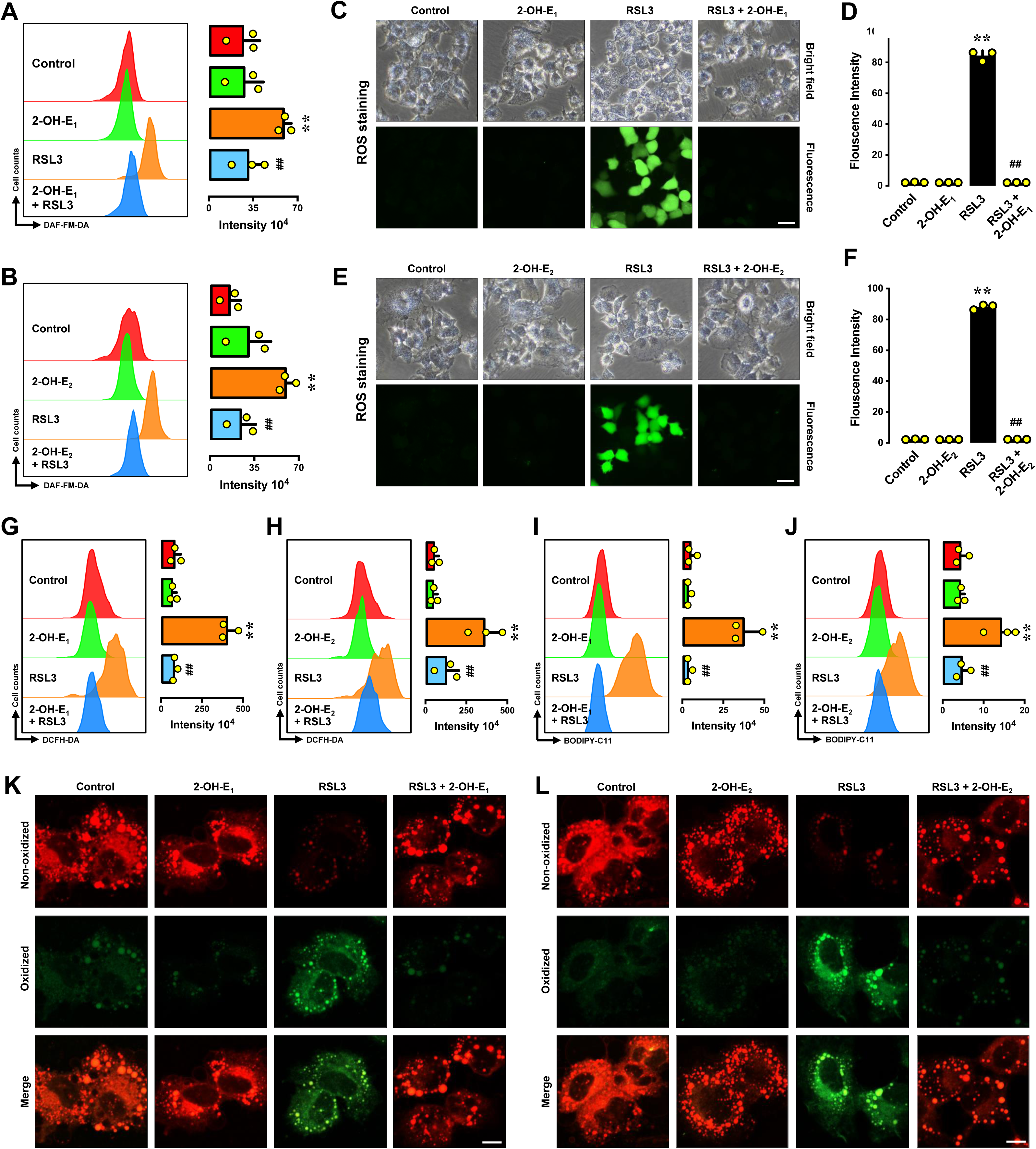
2-OH-E_1_ and 2-OH-E_2_ abrogate RSL3-induced accumulation of cellular NO, ROS and lipid-ROS in HuH-7 human hepatoma cells. **A, B.** Cellular level of NO after 4-h treatment with RSL3 (0.5 μM) ± 2-OH-E_1_ (10 μM) or 2-OH-E_2_ (10 μM). The left panels of **A** and **B** are flow cytometry data, and their right panels are the respective quantitative intensity values (n = 3). **C–H.** Cellular level of ROS after 4-h treatment with RSL3 (0.5 μM) ± 2-OH-E_1_ (10 μM) or 2-OH-E_2_ (10 μM). **C, D** are fluorescence microscopy images (scale bar = 60 μm), and **D**, **F** are respective quantitative intensity values (n = 3). **G, H.** The left panels are flow cytometry histograms, and the right panels are the respective quantitative intensity values (n = 3). **I–L.** Cellular level of lipid-ROS after 4-h treatment with RSL3 (0.5 μM) ± 2-OH-E_1_ (10 μM) or 2-OH-E_2_ (10 μM). The left panels of **I** and **J** are flow cytometry data, and their right panels are the respective quantitative intensity values (n = 3). **K** and **L** are confocal microscopy images (scale bar = 20 μm). All quantitative data are presented as mean ± S.D. (* or ^#^ *P* < 0.05; ** or ^##^ *P* < 0.01).

Together, these results demonstrate that both 2-OH-E_1_ and 2-OH-E_2_ can strongly abrogate erastin/RSL3-induced accumulation of intracellular NO, ROS and lipid-ROS in both H-4-II-E and HuH-7 hepatoma cells, and these effects are believed to contribute importantly to the protection against chemically-Induced ferroptotic cell death.

### 2-OH-E_1_ and 2-OH-E_2_ can bind directly to PDI

The above observations prompted us to identify the cellular target(s) that mediates the cytoprotective effect of 2-OH-E_1_ and 2-OH-E_2_. Our recent studies have shown, for the first time, that PDI is a cellular protein that mediates chemically-induced ferroptotic cell death (24). Next, we sought to determine whether PDI is a cellular target that also mediates the protective effect of 2-OH-E_1_ and 2-OH-E_2_ against chemically-induced ferroptosis in H-4-II-E cells and HuH-7 cells.

First, we determined whether 2-OH-E_1_ and 2-OH-E_2_ can bind to PDI by using the surface plasmon resonance assay. We found that 2-OH-E_1_ can bind to PDI with a high binding affinity (*K*_d_ = 24.2 and 27.9 nM, respectively, based on curve fitting and Scatchard plotting, respectively, **Fig. 7A, 7B**). In comparison, the apparent binding affinity of 2-OH-E_2_ for PDI is lower (*K*_d_ = 61.3–68.3 nM, **Fig. 7C, 7D**).

**Figure 7.**
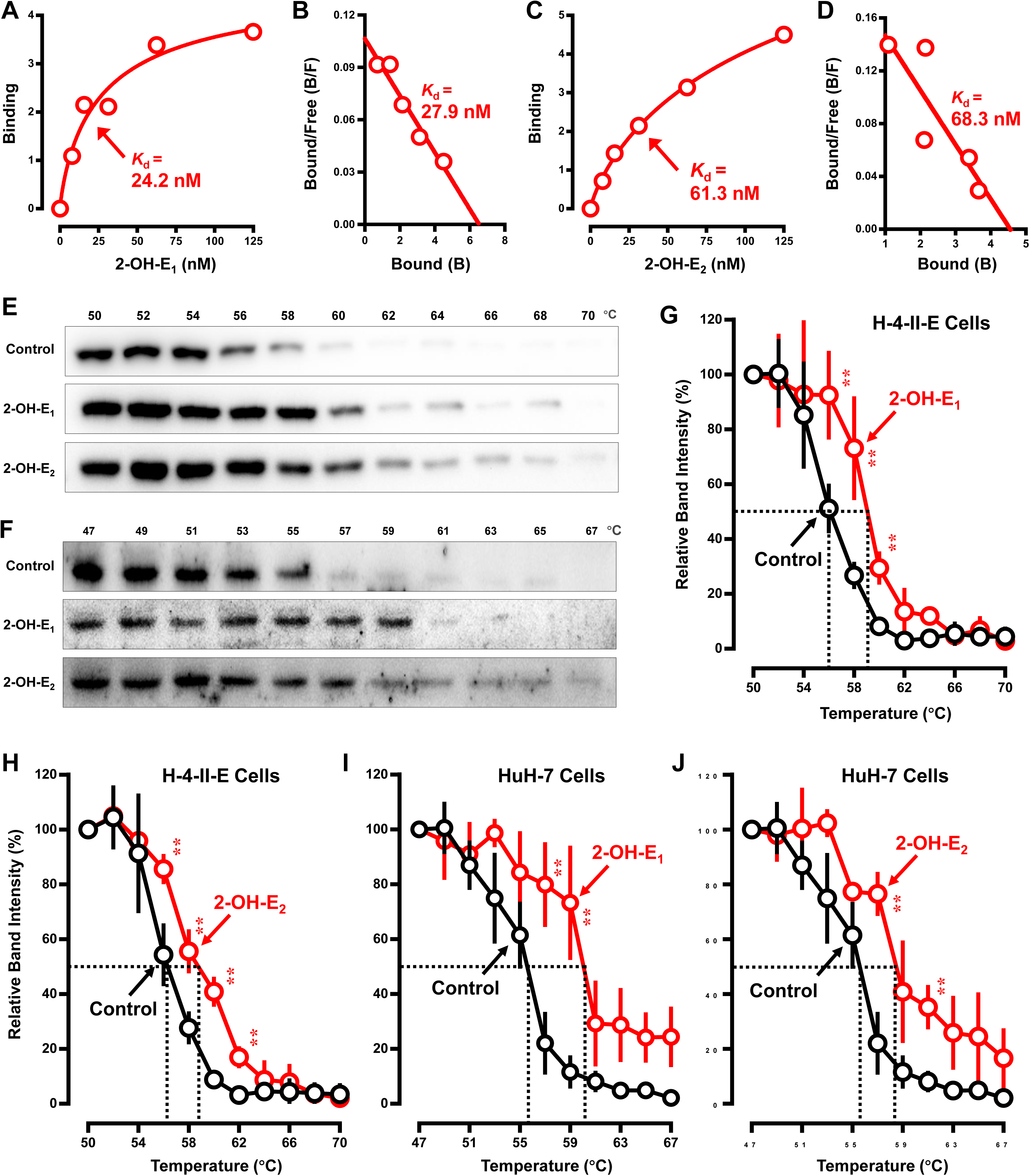
2-OH-E_1_ and 2-OH-E_2_ can bind directly to PDI. **A–D.** Surface plasmon resonance analysis of the binding affinity of 2-OH-E_1_ (**A** and **B**) and 2-OH-E_2_ (**C** and **D**) for purified PDI protein, showing the concentration-dependent binding curve (**A** and **C**) and the corresponding Scatchard plots (**B** and **D**). **E–J.** CETSA of live H-4-II-E cells (**E, G, H**) or live HuH-7 cells (**F, I, J**) treated with or without 2-OH-E_1_ or 2-OH-E_2_ (n = 3). The intensity of the protein bands was quantified using the Image J software. For CETSA curves, the relative band intensity was individually calculated relative to the protein intensity at the lowest temperature for the catechol estrogen-treated sample and the control samples, respectively. All quantitative data are presented as mean ± S.D. (* *P* < 0.05; ** *P* < 0.01).

Next, we performed the cellular thermal shift assay (CETSA) to evaluate the ability of these two catechol estrogens to bind to PDI protein in live H-4-II-E and HuH-7 cells. Based on Western blot analysis of PDI protein stability, a thermal shift associated with PDI protein was observed in catechol estrogen-treated cells compared to the control cells (**Fig. 7E, 7F**). A change in the Tm_50_ values (*i.e.*, the temperature at which 50% of the protein is precipitated by thermal denaturation) was determined to reflect the direct binding interaction of PDI protein with 2-OH-E_1_ or 2-OH-E_2_ in cultured cells. PDI had a Tm_50_ value of 56°C in control H-4-II-E cells (in the absence of a catechol estrogen) (**Fig. 7E, 7G**), but the presence of 2-OH-E_1_ or 2-OH-E_2_ increased its Tm_50_ to 59°C and 58.5°C, respectively, resulting in an increase in the Tm_50_ values (*i.e.*, *Δ*Tm_50_) by 3 and 2.5°C (**Fig. 7G, 7H**). A similarly thermal shift associated with PDI protein was also observed in HuH-7 cells jointly treated with 2-OH-E_1_ or 2-OH-E_2_ compared to the control cells (**Fig. 7F, 7I, 7J**). PDI in HuH-7 cells had a Tm_50_ value of 55.5°C (**Fig. 7I**), but the presence of 2-OH-E_1_ or 2-OH-E_2_ increased its Tm_50_ to 60 and 58.5°C, respectively, thus increasing the Tm_50_ values (*i.e.*, *Δ*Tm_50_) by 4.5 and 3°C (**Fig. 7I, 7J**). The CETSA results suggest that 2-OH-E_1_ has a higher binding affinity for PDI protein than does 2-OH-E_2_, which is also in agreement with the results of the surface plasmon resonance assay.

Computational molecular docking analysis was used to help understand how 2-OH-E_1_ and 2-OH-E_2_ bind to the PDI protein. The representative predicted structures of PDI–2-OH-E_1_ and PDI–2-OH-E_2_ complexes are shown in **Fig. 8A** and **8B**. Docking results reveal that 2-OH-E_1_ and 2-OH-E_2_ can tightly interact with PDI’s b’ domain and form a hydrogen bond with its His256 residue (**Fig. 8A, 8B**). When PDI’s His256 is mutated to Ala256, the hydrogen bond formed between His256 and the catechol estrogen (2-OH-E_1_ or 2-OH-E_2_) disappears (compare **Fig. 8C, 8E** with **Fig. 8I, 8K**). The surfaces of the binding pockets are shown in **Fig. 8D, 8F, 8J, 8L**). The difference in the binding pockets of the wild-type and mutant PDI proteins mostly centered around His256 which is mutated from a basic amino acid to a hydrophobic amino acid. The predicted binding affinities based on the representative docking structures of the wild-type PDI-His256–2-OH-E_1/2_ complexes are summarized in **Table 1**. The binding energies are increased after the His11Ala mutation, indicating that the stability of the interactions between PDI and 2-OH-E_1_ or 2-OH-E_2_ is decreased by the mutation. The modeling results provide partial support for the suggestion that PDI-His256 plays an important role in its binding interaction with 2-OH-E_1_ and 2-OH-E_2_. In addition, the binding energy values are also in line with the observations that 2-OH-E_1_ has a slightly higher binding affinity for the wild-type PDI protein than does 2-OH-E_2_.

**Figure 8.**
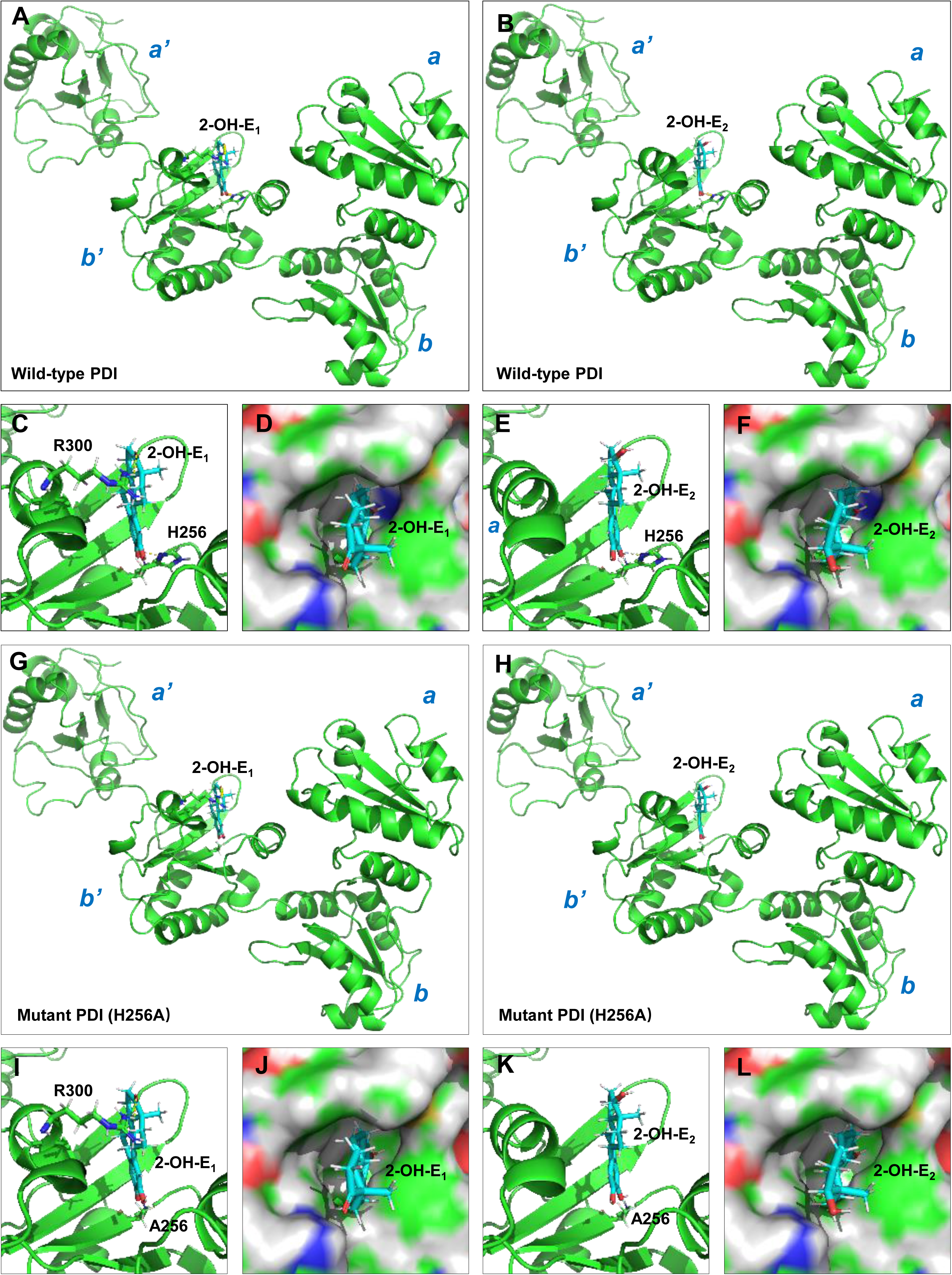
Predicted structures of the PDI–ligand complexes by molecular docking analysis. **A–B.** Structures of the binding complexes between the wild-type PDI and 2-OH-E_1_ (**A**) or 2-OH-E_2_ (**B**). **C, D, E, F.** Hydrogen bonds formed between PDI and 2-OH-E_1_ or 2-OH-E_2_ (**C, E**) and the surfaces of the binding pockets (**D, F**). PDI and 2-OH-E_1/2_ are colored in green and cyan, respectively. **G, H.** Structures of the binding complexes between the mutant PDI-Ala256 and 2-OH-E_1_ (**G**) or 2-OH-E_2_ (**H**). **I, J, K, L.** Hydrogen bonds formed between the mutant PDI and 2-OH-E_1_ or 2-OH-E_2_ (**I, K**) and the surfaces of the binding pockets (**J, L**). PDI and 2-OH-E_1/2_ are colored in green and cyan, respectively.

**Table 1.** Predicted binding affinities for the representative structures of the wild-type PDI–2-OH-E_1/2_ complexes and the mutant PDI–2-OH-E_1/2_ complexes obtained from molecular dynamics simulations.

| Protein | Ligand | Number of H-bonds between ligand and PDI protein | Number of H-bonds between ligand and PDI's residue-256 | PRODIDY-LIG (Kcal/mol) |
| --- | --- | --- | --- | --- |
| PDI-His256 | 2-OH-E<sub>1</sub> | 2 | 1 | -7.79 |
| PDI-Ala256 | 2-OH-E<sub>1</sub> | 1 | 0 | -7.6 |
| PDI-His256 | 2-OH-E<sub>2</sub> | 1 | 1 | -7.78 |
| PDI-Ala256 | 2-OH-E<sub>2</sub> | 0 | 0 | -7.59 |

MD simulations were carried out to further compare the binding affinities of the wild-type PDI–2-OH-E_1/2_ complexes. After 100-ns MD simulations, 2-OH-E_1_ and 2-OH-E_2_ are still bound inside the original binding sites (**Fig. 9A, 9B**). The predicted binding energy (according to PRODIGY-LIG (29)) for the wild-type PDI–2-OH-E_1_ complex is lower than that for the wild-type PDI–2-OH-E_2_ complex after 20-ns’ simulation (**Fig. 9C**), which indicates that the binding affinity between the wild-type PDI and 2-OH-E_1_ is higher than the binding affinity between the wild-type PDI and 2-OH-E_2_.

**Figure 9.**
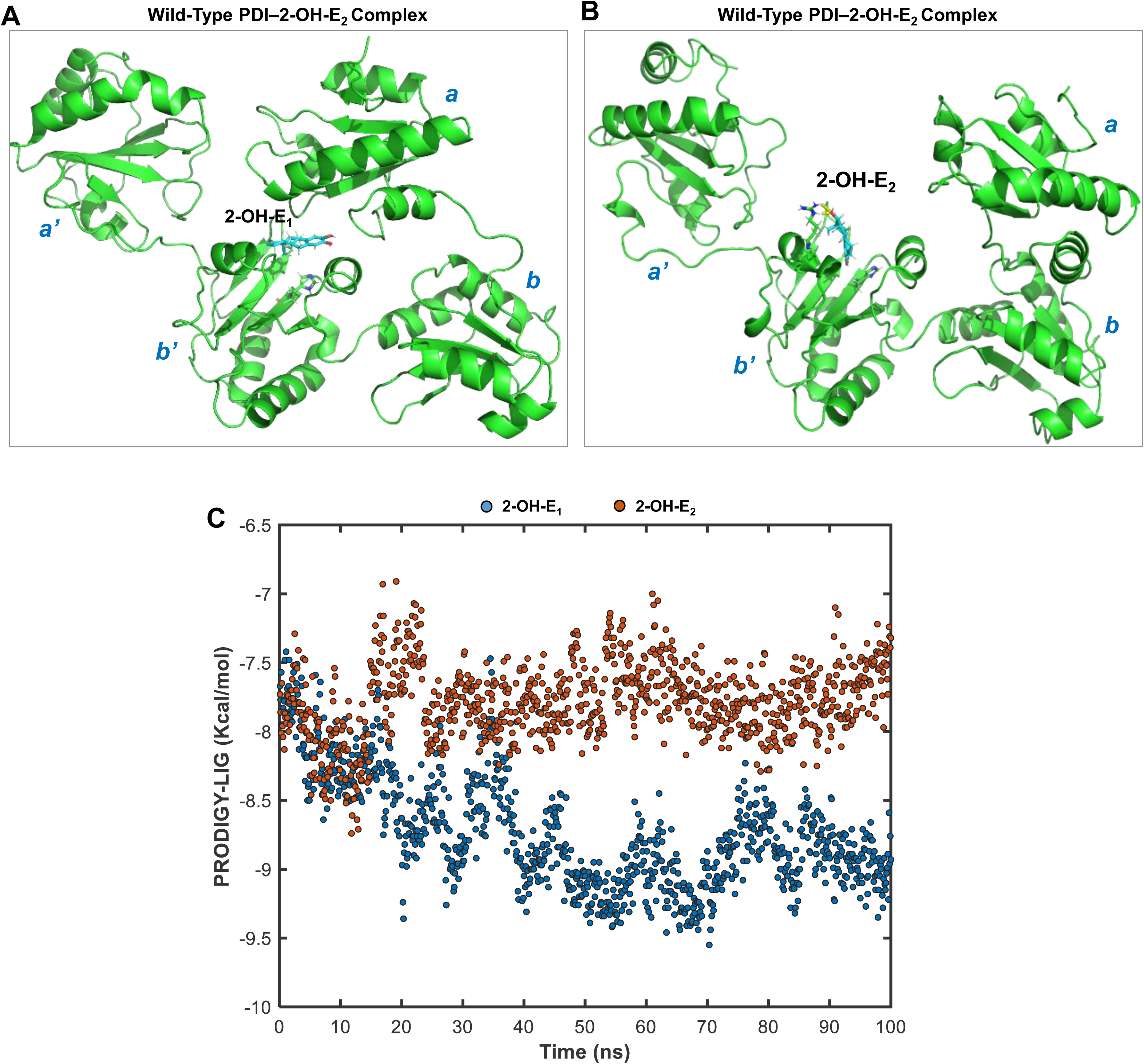
Conformations and binding affinity of the wild-type PDI–2-OH-E_½_ complexes obtained after MD simulation. **A.** Conformation of the wild-type PDI–2-OH-E_1_ complex. **B.** Confirmation of the wild-type PDI–2-OH-E_2_ complex. **C.** Computed binding energy values (PRODIGY-LIG scores) of the wild-type PDI–2-OH-E_1/2_ binding complexes during the MD simulation process.

### 2-OH-E_1_ and 2-OH-E_2_ can inhibit PDI’s catalytic activity

Results from our recent study (16) showed that the disulfide isomerase activity of PDI (which catalyzes disulfide bond formation in cellular protein substrates) mediates chemically-induced ferroptosis. Next, we sought to determine whether the ability of 2-OH-E_1_ and 2-OH-E_2_ to bind to PDI can result in inhibition of PDI’s catalytic activity by using the *in vitro* insulin aggregation assay. Insulin has two chains cross-linked together by disulfide bonds, and PDI can break these disulfide bonds to facilitate the formation of aggregates, which has been commonly used to reflect PDI’s catalytic activity (41). We found that PDI’s catalytic activity *in vitro* was suppressed by the presence of 2-OH-E_1_ or 2-OH-E_2_ in a concentration-dependent manner (**Fig. 10A, 10B**).

**Figure 10.**
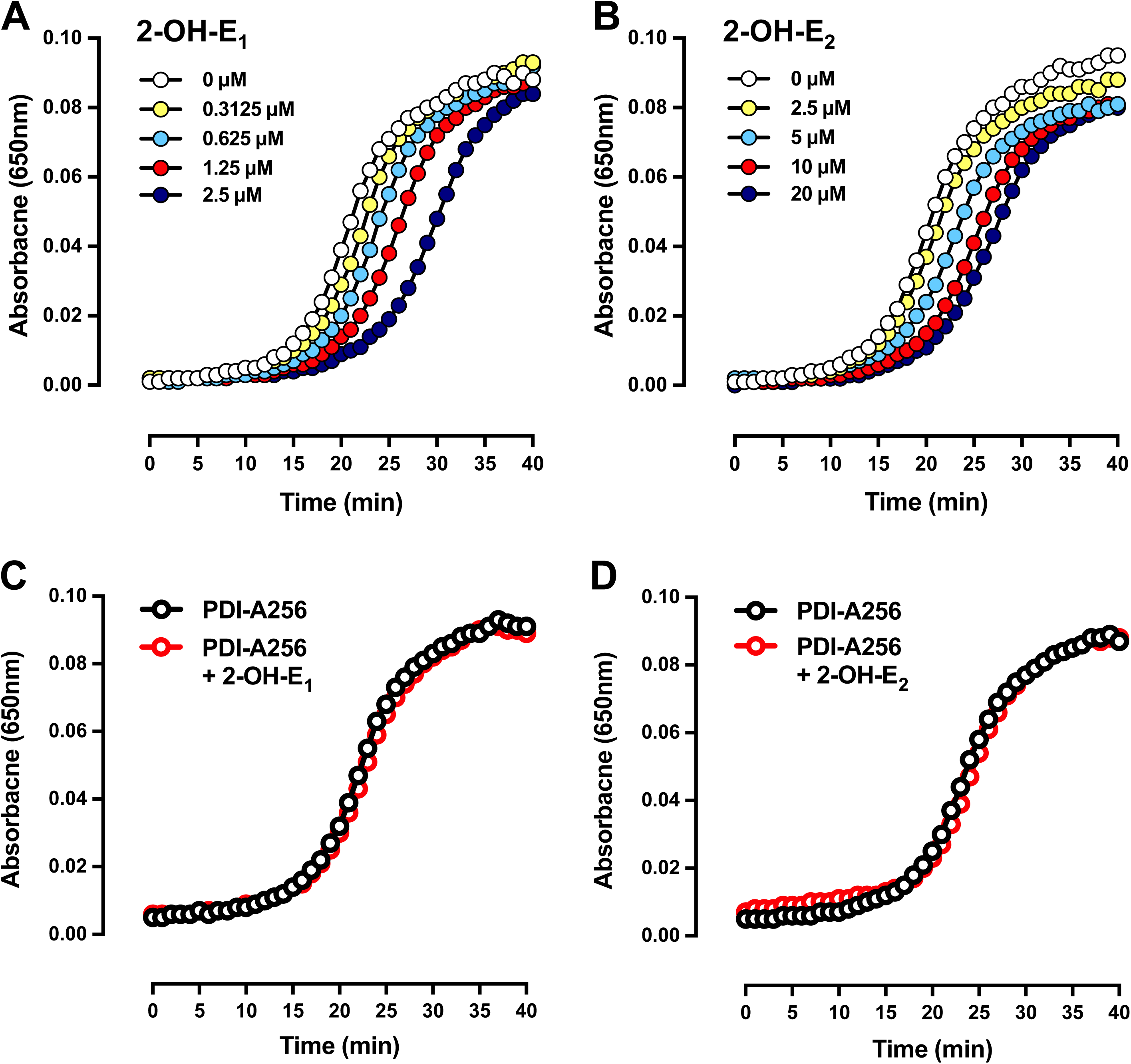
2-OH-E_1_ and 2-OH-E_2_ inhibit the catalytic activity of the wild-type PDI but not the mutant PDI-Ala256. **A, B, C, D.** Inhibitory effect of 2-OH-E_1_ and 2-OH-E_2_ on the catalytic activity of the wild-type PDI (**A, B**) and mutant PDI-Ala256 (**C, D**). PDI catalytic activity was measured by determining PDI-mediated insulin aggregation. Each experiment was repeated at least three times with similar results.

To provide experimental support for docking prediction that PDI’s His256 is important for its binding interaction with 2-OH-E_1_ or 2-OH-E_2_, next we chose to compare the ability of 2-OH-E_1_ and 2-OH-E_2_ to inhibit the catalytic activity of the wild-type and mutant PDIs. While the mutant PDI-Ala256 had a slightly-reduced catalytic velocity (*i.e.*, it would take longer to catalyze the same reaction), its catalytic ability to producing the same maximal level of insulin aggregates was not diminished (**Fig. 10A, 10B**). Notably, the presence of 2-OH-E_1_ or 2-OH-E_2_ no longer showed a meaningful inhibition of PDI-Ala256’s catalytic activity in mediating insulin aggregation (**Fig. 10C, 10D**). This result suggested that 2-OH-E_1_ and 2-OH-E_2_ could not effectively bind to the mutant PDI-Ala256 to exert the same degree of inhibition of its catalytic activity as it did with the wild-type PDI protein. Together, the results from the *in vitro* enzymatic assay demonstrate that while 2-OH-E_1_ and 2-OH-E_2_ can directly bind to the wild-type human PDI and inhibit its catalytic activity, they cannot do so with the mutant PDI-Ala256.

### 2-OH-E_1_ and 2-OH-E_2_ can abrogate erastin/RSL3-induced iNOS upregulation and its dimerization

Our recent studies reported that during erastin– and RSL3-induced ferroptosis, there is an induction of cellular NOS protein level (14,16,40). In this study, therefore, we sought to determine whether 2-OH-E_1_ and 2-OH-E_2_ affect iNOS induction and dimer formation in cultured hepatoma cells following exposure to erastin and/or RSL3. In H-4-II-E cells, treatment with erastin alone increased cellular levels of iNOS protein and its dimer compared to the control groups (**Fig. 11A, 11B**). While treatment of cells with 10 μM 2-OH-E_1_ (**Fig. 11A**) or 2-OH-E_2_ (**Fig. 11B**) alone did not significantly affect the cellular levels of iNOS protein and its dimer, joint treatment of the cells with erastin + 2-OH-E_1_ (**Fig. 11A**) or 2-OH-E_2_ (**Fig. 11B**) effectively abrogated erastin-induced iNOS protein upregulation and dimer formation. In comparison, the PDI protein level remained largely unchanged by these treatments (**Fig. 11A, 11B**).

**Figure 11.**
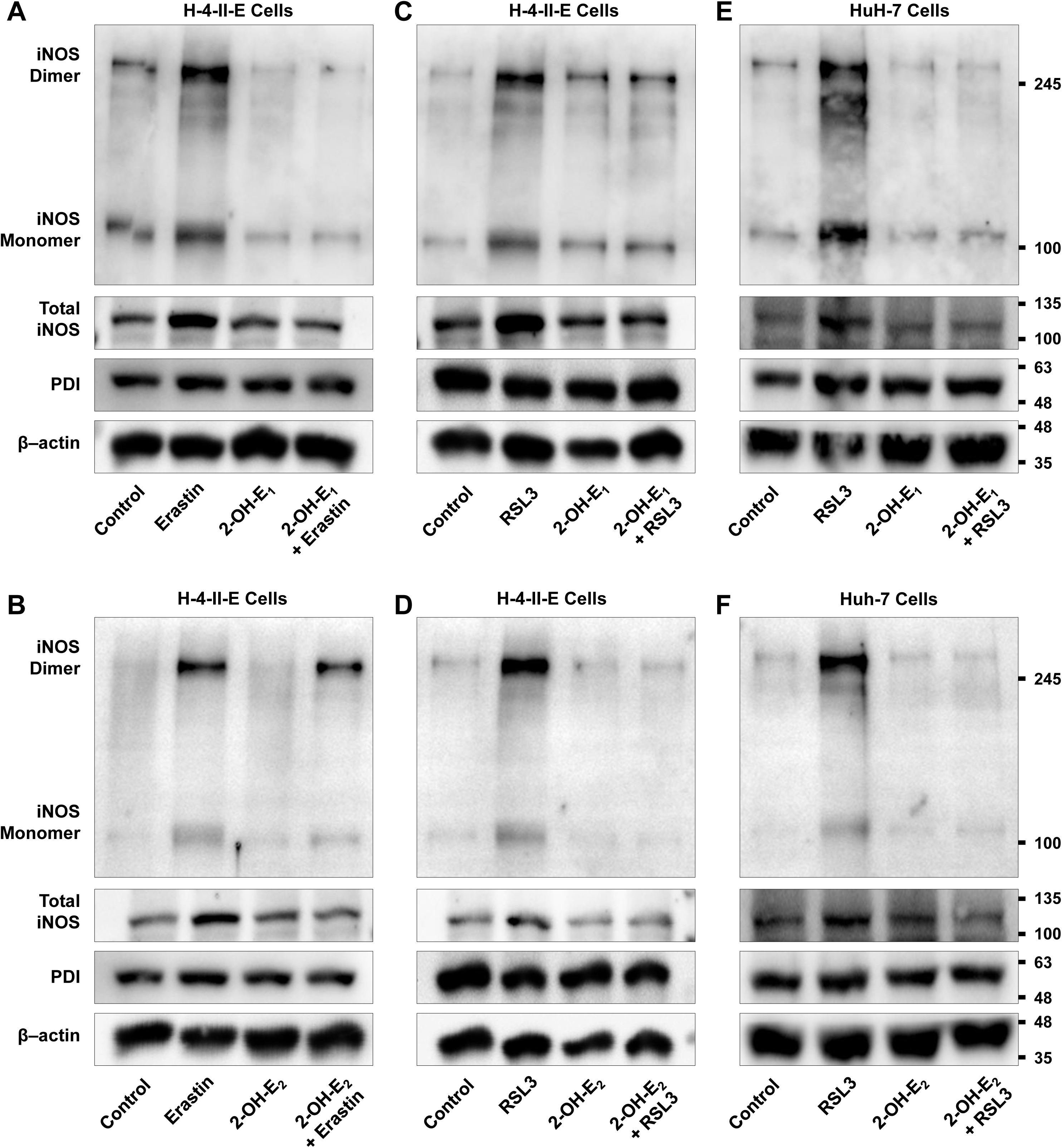
2-OH-E_1_ and 2-OH-E_2_ abrogate erastin/RSL3-induced upregulation of iNOS protein level and its dimerization. **A, B.** Level of total iNOS protein and its monomer and dimer following 8-h treatment with 1 μM erastin ± 7.5 μM 2-OH-E_1_ (**A**) or 5 μM 2-OH-E_2_ (**B**) in H-4-II-E rat hepatoma cells. The total cellular proteins (25 μg) were separated by 8% SDS-PAGE under reducing conditions or by 6% SDS-PAGE under non-reducing conditions, and then immunoblotted with specific antibodies for iNOS, PDI and β-actin. **C, D.** Level of total iNOS protein and its monomer and dimer following 4-h treatment with 0.4 μM RSL3 ± 7.5 μM 2-OH-E_1_ (**C**) or 5 μM 2-OH-E_2_ (**D**) in H-4-II-E rat hepatoma cells. The total cellular proteins (25 μg) were separated by 8% SDS-PAGE under reducing conditions or by 6% SDS-PAGE under non-reducing conditions, and then immunoblotted with specific antibodies for iNOS, PDI and β-actin. **E, F.** Level of total iNOS protein and its monomer and dimer following 2-h treatment with 0.5 μM RSL3 ± 10 μM 2-OH-E_1_ (**E**) or 10 μM 2-OH-E_2_ (**F**) in HuH-7 human hepatoma cells. The total cellular proteins (25 μg) were separated by 8% SDS-PAGE under reducing conditions or 6% SDS-PAGE under non-reducing conditions, and then immunoblotted with specific antibodies for iNOS, PDI and β-actin.

Very similar observations on cellular iNOS, its dimerization and PDI levels were made in H-4-II-E cells when they were treated with RSL3 ± 2-OH-E_1_ or 2-OH-E_2_ (**Fig. 11C, 11D**). Treatment with RSL3 alone increased iNOS protein level and its dimerization, and joint treatment of cells with 10 μM 2-OH-E_1_ or 2-OH-E_2_ abrogated RSL3-induced iNOS upregulation and dimer formation. Cellular PDI level was not significantly affected by these treatments (**Fig. 11C, 11D**).

For comparison, we have also determined the effect of 2-OH-E_1_ and 2-OH-E_2_ on RSL3-induced iNOS upregulation and dimer formation in HuH-7 cells. Treatment of these cells with RSL3 alone caused an increase in iNOS protein level compared to the control, but joint treatment of cells with RSL3 + 2-OH-E_1_ (**Fig. 11E**) or 2-OH-E_2_ (**Fig. 11F**) effectively abrogated RSL3-induced iNOS upregulation and dimer formation. In comparison, PDI protein levels remained mostly unchanged by these treatments. Together, these results indicate that treatment of cells with 2-OH-E_1_ and 2-OH-E_2_ significantly abrogates chemically-induced upregulation of cellular iNOS protein and its dimerization.

To provide additional support for the hypothesis that PDI–iNOS inhibition plays a major role in mediating the cytoprotective action of 2-OH-E_1_ and 2-OH-E_2_, we have also tested, for comparison, the effect of cystamine (a known PDI inhibitor (41)), SMT (a known iNOS inhibitor (42)) and cPTIO (an NO scavenger (43)) on erastin-induced ferroptotic cell death and the accumulation of cellular NO, ROS and lipid-ROS in H-4-II-E cells (a representative cell line tested in this study). We found that joint treatment of the cells with cystamine (which does not have any reducing ability) strongly protected the cells against erastin-induced cell death (**Fig. 12A**), accompanied by strong abrogation of erastin-induced accumulation of cellular NO, ROS and lipid-ROS (**Fig. 12D –12F**). Similarly, joint treatment of the cells with SMT also strongly protected the cells against erastin-induced cell death (**Fig. 12B**), together with strong abrogation of erastin-induced accumulation of cellular NO, ROS and lipid-ROS (**Fig. 12G –12I**). Lastly, joint treatment of the cells with cPTIO elicited a partial protection against erastin-induced cell death (**Fig. 12C**), which is also accompanied by a partial reduction in erastin-induced accumulation of cellular NO, ROS and lipid-ROS (**Fig. 12J –12L**). Together, these results provide additional support for the notion that pharmacological inhibition of PDI or iNOS/NO function can effectively abrogate erastin-induced ferroptotic cell death.

**Figure 12.**
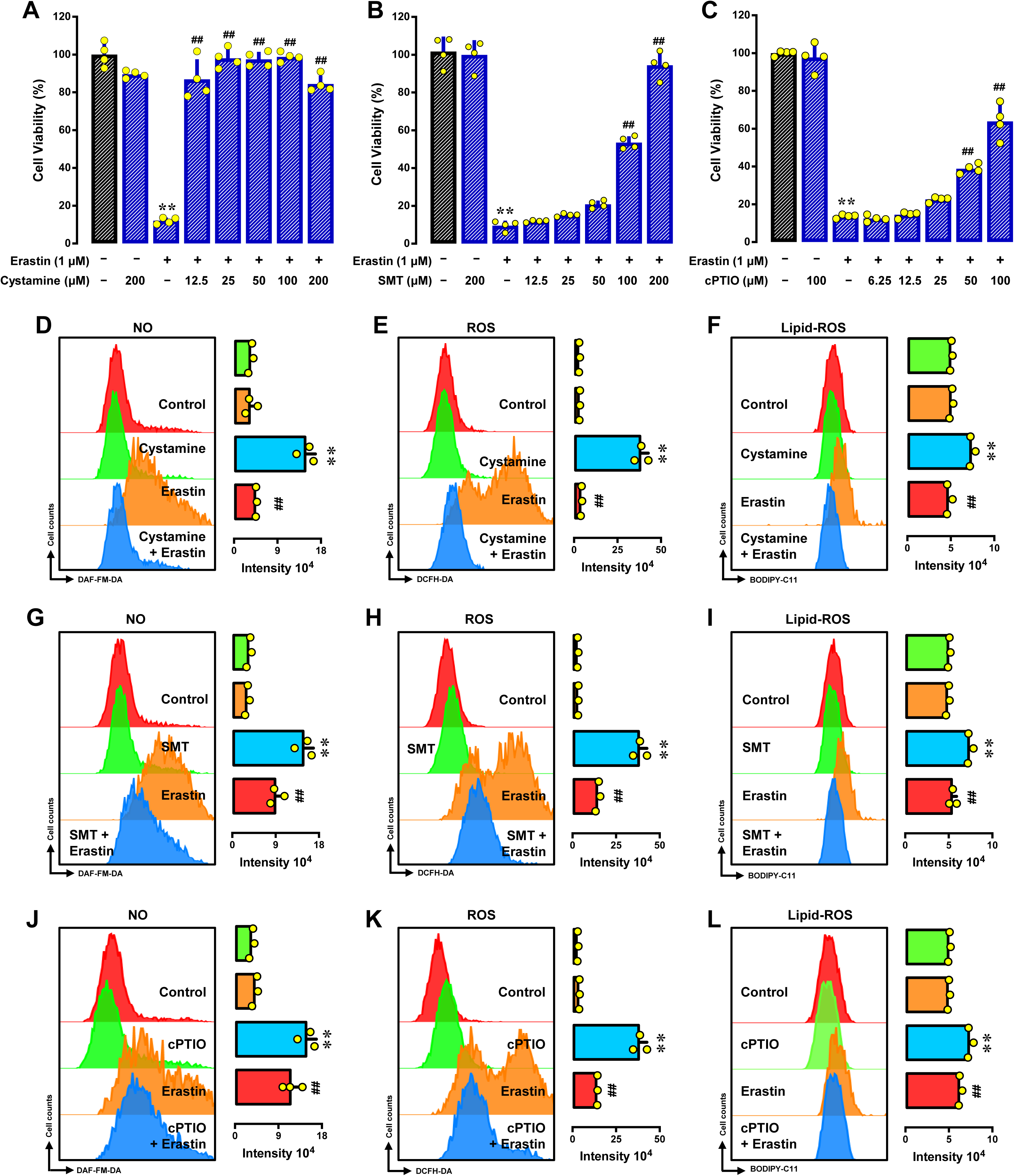
Effect of cystamine, SMT and cPTIO on erastin-induced cell death and cellular accumulation of NO, ROS and lipid-ROS in H4-II-E cells. **A, B, C.** Cells were treated with erastin (1 μM) ± cystamine (12.5□200 μM) or SMT (12.5□200 μM) or cPTIO (6.25□100 μM) for 24 h, and cell viability was then determined by MTT assay (n = 4). **D**LJ**L.** Flow cytometry analysis of cellular levels of NO, ROS and lipid-ROS after 8-h treatment with erastin (1 μM) ± cystamine (100 μM; **D–F**) or SMT (200 μM; **G–I**) or cPTIO (100 μM; **J–L**). Each left panel is the flow cytometry data, and each right panel is the respective quantitative intensity values (n = 3). All quantitative data are presented as mean ± S.D. (* or ^#^ *P* < 0.05; ** or ^##^ *P* < 0.01).

### 2-OH-E_1_ and 2-OH-E_2_ protect against APAP-induced liver injury in vivo

#### Gross and histological change

APAP is commonly used to induce liver injury in animal models through induction of GSH depletion-associated hepatocyte ferroptosis (44). In this study, APAP-induced acute liver injury in male C57BL/6J mice was used as an *in vivo* model to investigate the hepatoprotective effect of E_1_, E_2_, 2-OH-E_1_ and 2-OH-E_2_. The mice were treated with APAP (300 mg/kg, gastric intubation) alone or in combination with an estrogen.

Administration of APAP alone for 24 h caused marked changes in liver surface abnormalities and in liver-to-body weight ratio, and joint treatment of the animals with 2-OH-E_1_ or 2-OH-E_2_ exerted a strong protection against APAP-induced acute liver injury based on gross changes in liver surface abnormality (**Fig. 13A**) and liver-to-body weight ratio (**Supplementary Fig. S1**). Surprisingly, E_1_ or E_2_ was similarly effective in protecting against APAP-induced liver injury *in vivo* compared to 2-OH-E_1_ or 2-OH-E_2_, respectively (**Fig. 13A**). Analysis of H/E-stained liver slides showed that APAP treatment caused extensive hepatocyte death, and severe hepatocyte degeneration was clearly seen around the central vein region of hepatic lobules. Typical cellular damage seen under a light microscope included cellular swollen and ballooning, loss of nuclei, and inflammatory cell infiltration (**Fig. 13B**). Joint treatment of the animals with E_1_, E_2_, 2-OH-E_1_ or 2-OH-E_2_ exerted a dose-dependent protection against APAP-induced hepatocyte damage, and the protection against APAP hepatotoxicity is more pronounced around the central vein region (**Fig. 13B**).

**Figure 13.**
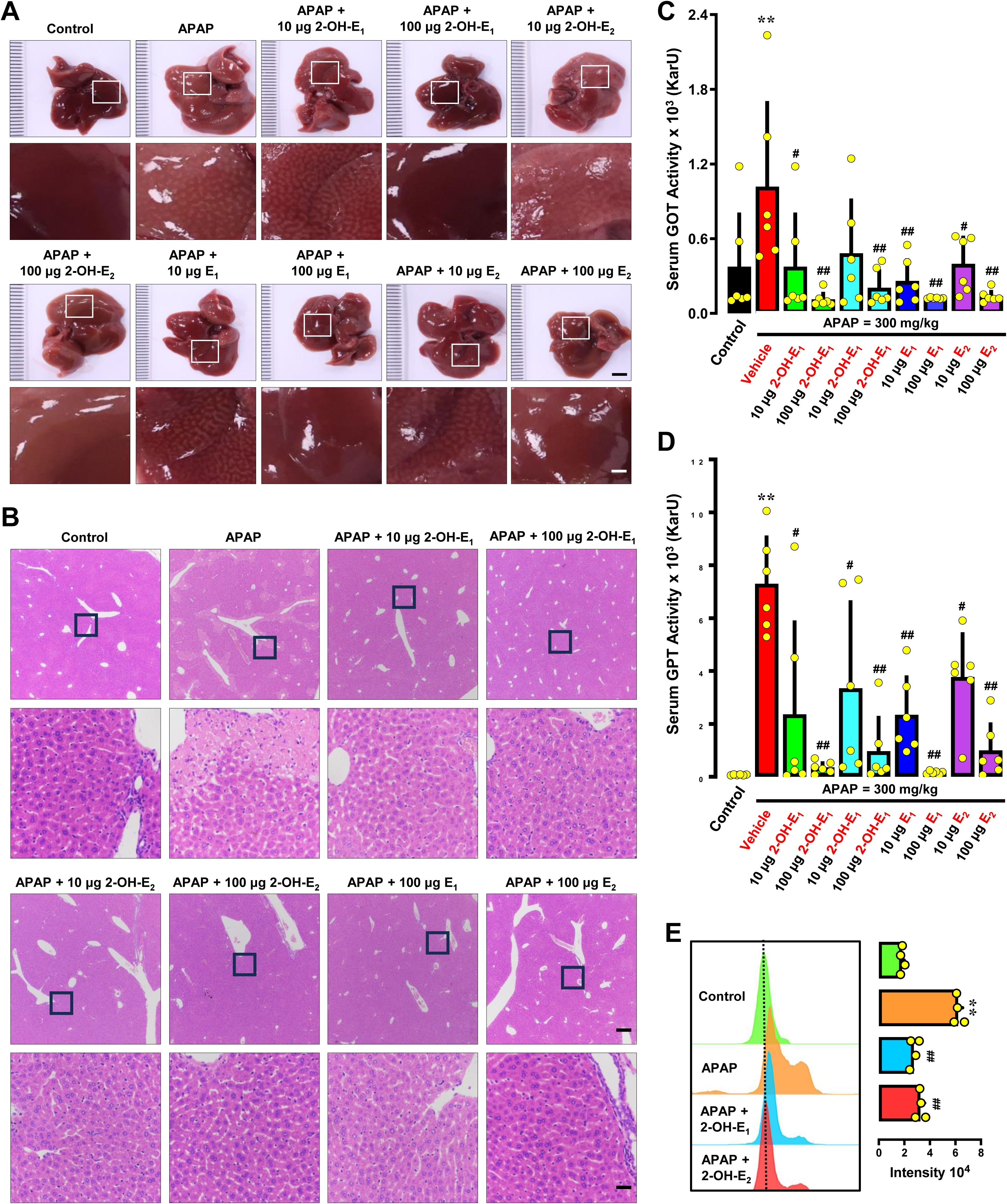
Role of PDI in APAP-induced liver injury in C57BL/6J mice. **A, B, C, D, E**. Protective effect of 2-OH-E_1_, 2-OH-E_2_, E_1_ and E_2_ against APAP-induced hepatocyte injury in C57BL/6J mice. After fasting for 12 h, the male C57BL/6J mice were treated with APAP (300 mg/kg, gastric intubation) ± each estrogen (10 and 100 μg/mouse, *i.p.*) for an additional 24 h. **A.** Lver gross morphology (scale bar = 4mm) and the enlarged region (scale bar = 1 mm). **B**. H/E staining (scale bar = 500 μm) and the enlarged region (scale bar = 50 μm). **C**. Plasma GOT levels (n = 6). **D.** Plasma GPT levels (n = 6). **E.** Hepatic ROS levels. Fresh liver samples (approximately 20 mg) were taken and weighed, stained with DCFH-DA (20 μM) for 50 min, and then analyzed by flow cytometry and the right panel is the quantitative intensity value (n = 4). All quantitative data are presented as mean ± S.D. (* or ^#^ *P* < 0.05; ** or ^##^ *P* < 0.01).

**Change in biochemical markers.**

The plasma GOT and GPT levels were commonly used to reflect the degree of liver injury following APAP treatment. APAP caused drastic increases in plasma GOT and GPT levels (∼1000– and 10-fold higher, respectively; *P* < 0.05) over the control group (**Fig. 13C, 13D**). Joint treatment of the animals with 2-OH-E_1_, 2-OH-E_2_, E_1_ or E_2_, significantly reduced the GOT and GPT levels (the GOT activity was nearly reduced to control level).

To probe the role of cellular ROS in APAP-induced hepatocyte toxicity *in vivo*, fresh mouse liver tissues were harvested for preparation of liver cell suspension. The isolated liver cells were stained for ROS and analyzed by flow cytometry. While 24-h APAP treatment induced ROS accumulation in mouse liver cells *in vivo*, joint treatment with 2-OH-E_1_, 2-OH-E_2_ E_1_ or E_2_ abrogated APAP-induced ROS accumulation in these cells (**Fig. 13E**).

## DISCUSSION

### 2-OH-E_1_ and 2-OH-E_2_ protect against chemically-induced ferroptosis

Ferroptosis is an iron-dependent form of regulated cell death characterized by the accumulation of lipid-ROS and subsequent oxidative damage. Ferroptosis has emerged as a distinct type of cell death with implications in various pathological conditions, including neurodegenerative diseases, ischemia-reperfusion injury, and chemically-induced, GSH depletion-associated organ damage. In recent years, there has been a growing interest in identifying potential therapeutic strategies to modulate ferroptosis and protect cells from its detrimental effect.

We demonstrate in this study that 2-OH-E_1_ and 2-OH-E_2_, two quantitatively-major catechol estrogen metabolites formed in the liver (18), have a strong protective effect against erastin-induced ferroptotic cell death in H-4-II-E and HuH-7 hepatoma cells *in vitro*. In addition, a significant NO accumulation was observed following treatment of these cells with erastin alone, and joint treatment with 2-OH-E_1_ or 2-OH-E_2_ effectively abrogates erastin-induced NO buildup. Similarly, erastin treatment also leads to significant accumulation of cellular ROS and lipid-ROS, and joint treatment with 2-OH-E_1_ or 2-OH-E_2_ abrogates erastin-induced accumulation of cellular ROS and lipid-ROS. Together, these observations demonstrate that 2-OH-E_1_ and 2-OH-E_2_ can effectively abrogate erastin-induced accumulation of cellular NO, ROS and lipid-ROS, which jointly contributes to the protection against erastin-induced ferroptotic cell death.

In this study, the protective effect of 2-OH-E_1_ and 2-OH-E_2_ against RSL3-induced ferroptosis in H-4-II-E and HuH-7 hepatoma cells is also investigated. RSL3 induces ferrotosis through its inhibition of GPX4, an enzyme involved in lipid-ROS detoxification (5,6). Inhibition of GPX4 activity leads to accumulation of lipid-ROS and subsequent cellular injury, ultimately culminating in ferroptotic cell death (5,6). Similar to the observations made with erastin, we also show that 2-OH-E_1_ and 2-OH-E_2_ elicit a strong dose-dependent protection against RSL3-induced ferroptosis in H-4-II-E and HuH-7 hepatoma cells. In addition, a similar abrogating effect of catechol estrogens on RSL3-induced accumulation of cellular NO, ROS and lipid-ROS is observed.

Experimental evidence is also provided in this study for the protective effect of 2-OH-E_1_ and 2-OH-E_2_ against APAP-induced, GSH depletion-associated liver injury in an *in vivo* mouse model. APAP is a commonly-used inducer of liver injury through the induction of GSH depletion-associated oxidative cell death in hepatocytes (44–46). We show that while administration of APAP alone for 24 h causes significant liver injury, joint treatment of animals with 2-OH-E_1_ or 2-OH-E_2_ elicits a dose-dependent protection against APAP-induced hepatocyte injury. In addition, treatment of mice with APAP for 24 h induces hepatocyte ROS accumulation *in vivo*, and joint treatment with 2-OH-E_1_ or 2-OH-E_2_ abrogates APAP-induced ROS accumulation in these cells. These observations made in a mouse model *in vivo* are in agreement with the observations made in cell culture experiments. Surprisingly, while E_1_ and E_2_ do not show a strong protective effect against chemically-induced ferroptosis in cultured hepatoma cells, they exert a similarly strong protective effect as 2-OH-E_1_ and 2-OH-E_2_ in the *in vivo* mosue model. Metabolic conversion of E_1_ and E_2_ to their respective 2-hydroxylated metabolites in the liver likely contributes importantly to their strong *in vivo* cytoprotective effect seen in this organ (discussed later).

In summary, the results of this study demonstrate that 2-OH-E_1_ and 2-OH-E_2_ have a strong protective effect against chemically-induced ferroptosis both *in vitro* and *in vivo*. In addition, E_1_ and E_2_ exert a similarly strong protective effect as 2-OH-E_1_ and 2-OH-E_2_ in an *in vivo* mouse model.

### PDI mediates the cytoprotective effect of 2-OH-E_1_ and 2-OH-E_2_

In this study, it is hypothesized that 2-OH-E_1_ and 2-OH-E_2_ can bind directly to cellular PDI protein and inhibit its catalytic activity, which then initiates downstream changes culminating in cytoprotection against ferroptosis. As summarized below, a series of experimental evidence are presented to offer support for the notion that 2-OH-E_1_ and 2-OH-E_2_ exert their protection against erastin/RSL3-induced ferroptosis by targeting PDI and inhibiting its catalytic activity, which then reduces PDI-mediated conversion of iNOS monomers to its catalytically-active dimers, and ultimately, prevents chemically-induced ferroptotic cell death.

i. Computational docking analysis indicates that 2-OH-E_1_ and 2-OH-E_2_ can bind tightly inside a rather deep binding pocket in the PDI protein, forming a hydrogen bond with His256.
ii. Surface plasmon resonance analysis shows that 2-OH-E_1_ and 2-OH-E_2_ can bind to purified PDI protein with a rather high binding affinity (apparent *K*_d_ of 24.21127.9 nM for 2-OH-E_1_ and 61.31168.3 nM for 2-OH-E_2_).
iii. Cellular thermal shift assay (CETSA) shows that 2-OH-E_1_ and 2-OH-E_2_ can bind to PDI in live hepatoma cells by causing an increase in the Tm_50_ value, demonstrating that 2-OH-E_1_ or 2-OH-E_2_ can bind to PDI protein in these cells.
iv. The *in vitro* enzymatic assay shows that 2-OH-E_1_ and 2-OH-E_2_ can directly inhibit PDI’s catalytic activity in a concentration-dependent manner, but this direct inhibition of PDI’s catalytic activity is not observed with the mutant PDI-Ala256 protein. Here it is of note that in the absence of inhibitors, the catalytic velocity of the mutant PDI-Ala256 protein is slightly slower than the wild-type PDI, but both PDI proteins still retain the same maximal catalytic ability (*i.e.*, the maximal insulin aggregation with both enzymes is nearly the same). Based on the information gained from computational modeling analysis, it is clear that the binding site of PDI for 2-OH-E_1_ or 2-OH-E_2_ is located between its b and b’ domains, whereas its catalytic site lies in the a and a’ domains which contain the CXXC sequences [10]. The reduced catalytic velocity of PDI-Ala256 and its insensitivity to inhibition by 2-OH-E_1_ and 2-OH-E_2_ are in agreement with the notion that His256 is involved in the binding interaction with both the protein substrates (such as insulin) and the catechol estrogen inhibitors. Lastly, since the mutant PDI-Ala256 protein still retains its maximal catalytic ability, this observation is also consistent with the fact that the catalytic site (which lies between the a and a**’** domains) is not affected by this mutation.

Together, these results demonstrate that 2-OH-E_1_ and 2-OH-E_2_ can directly bind to PDI, and the formation of a hydrogen bond with PDI-His256 contributes importantly to their binding interaction with PDI. It is clear that the binding interaction of 2-OH-E_1_ and 2-OH-E_2_ with PDI is associated with their inhibition of PDI’s catalytic activity.

In the liver, iNOS is an enzyme responsible for the production of NO in response to various stimuli (47). Recent studies from our laboratory have shown that treatment of cells with erastin or RSL3 can lead to PDI activation, and the activated PDI then catalyzes the conversion of the NOS monomer to its dimer form, which increases NO production (14,16,40). The elevation of NO levels subsequently leads to accumulation of cellular ROS and lipid-ROS, ultimately resulting in ferroptosis (14,16,82). The notion that 2-OH-E_1_ and 2-OH-E_2_ can bind to PDI and effectively inhibit its catalytic activity is also supported by the biochemical changes observed in erastin/RSL3-treated live hepatoma cells. Specifically, we find that treatment of the cells with erastin alone results in PDI-mediated iNOS activation (*i.e.*, formation of iNOS dimers) along with cellular NO accumulation, and joint treatment of the cells with 2-OH-E_1_ or 2-OH-E_2_ abrogates erastin-induced iNOS dimerization and NO accumulation. In addition, 2-OH-E_1_ and 2-OH-E_2_ also suppresses erastin-induced upregulation of cellular iNOS protein, which also contributes importantly to protection against erastin-induced cytotoxicity. Together, these results demonstrate that 2-OH-E_1_ and 2-OH-E_2_ effectively protect against erastin-induced ferroptotic cell death by abrogating iNOS upregulation and by targeting PDI-mediated iNOS dimerization and the subsequent accumulation of cellular NO and ROS/lipid-ROS.

The importance of the PDI□iNOS/NO□ROS/lipid-ROS axis as a target for protection against erastin-induced ferroptosis by 2-OH-E_1_ and 2-OH-E_2_ is further illustrated by additional analysis of the protective effect of cystamine, SMT and cPTIO. We find that joint treatment of cells with cystamine strongly abrogates erastin-induced accumulation of cellular NO and ROS/lipid-ROS, which is coupled with a strong protection against ferroptosis. The strong abrogation of cellular NO and ROS/lipid-ROS accumulation by cystamine is clearly not due to its direct free radical-scavenging activity as this chemical (NH_2_CH_2_CH_2_11S11S11CH_2_CH_2_NH_2_) is not a reducing agent; rather, it is a weak oxidizing agent. Its effect is purely due to its ability to covalently modify the free –SH group(s) in PDI’s catalytic site, resulting in the formation of PDI–S11S11CH_2_CH_2_NH_2_. Similarly, observations on the protective effect of SMT and cPTIO offer additional support for the crucial role of iNOS and NO in mediating erastin-induced ferroptosis. Here it is of note that the protective effect of cPTIO is rather modest (weaker than the effect of cystamine and SMT), likely due to its relatively weaker NO-scavenging capacity at the nontoxic concentration used in this experiment. Jointly, these additional results further demonstrate that inhibition of the PDI□iNOS/NO axis is a major mechanism underlying the protective action of 2-OH-E_1_ and 2-OH-E_2_ against erastin-induced ferroptosis.

Based on the observations made in this study, it is evident that PDI is also involved in mediating RSL3-induced ferroptosis in hepatoma cells, and inhibition of PDI’s function by 2-OH-E_1_ and 2-OH-E_2_ contributes importantly to their protection against RSL3-induced cell death. It has been long held the view that the ferroptosis-inducing activity of RSL3 is primarily due to its inhibition of GPX4 (6). However, a recent study has reported that RSL3 can also strongly inhibit the enzymatic activity of thioredoxin reductase 1 (TrxR1) (48). Theoretically, inhibition of TrxR1 by RSL3 will shift the pool of cellular PDI protein (a member of the thioredoxin superfamily) toward the catalytically-active oxidized state, thus favoring iNOS dimer formation. Offering support this hypothesis, recently we have shown (40) that in addition to inhibition of GPX4, RSL3, through its ability to inhibit TrxR1 enzymatic activity (48), can promote ferroptosis by facilitating NOS dimerization, followed by cellular NO, ROS and lipid-ROS accumulation, and ultimately ferroptotic cell death (40). This novel mechanism of RSL3-induced ferroptosis also provides a mechanistic basis for the observed strong cytoprotective action of 2-OH-E_1_ and 2-OH-E_2_ against RSL3-induced ferroptosis.

Lastly, it is of note that besides inhibition of PDI-mediated iNOS activation by 2-OH-E_1_ and 2-OH-E_2_, joint treatment of cells with these two catechol estrogens also strongly reduces RSL3-induced upregulation of iNOS protein level, which is another factor contributing to their cytoprotective action. In addition, as 2-OH-E_1_ and 2-OH-E_2_ are phenolic compounds, their direct antioxidant property may also contribute, to certain degrees, to their overall protective effect against chemically-induced ferroptosis.

### E_1_ and E_2_ exert hepatocyte protection *in vivo*

2-Hydroxylation of E_2_ and E_1_ to 2-OH-E_2_ and 2-OH-E_1_, respectively, is a major metabolic pathway in the liver, whereas 4-hydroxylation to 4-OH-E_2_ and 4-OH-E_1_, respectively, represents a quantitatively-minor pathway (usually <15% of 2-hydroxylation) in this organ (18). Several different CYP isoforms can contribute, to varying degrees, to the 2-hydroxylation of E_2_ and E_1_ in the liver of rodents and humans (18). Both 2-OH-E_2_ and 2-OH-E_1_ can bind to the ERα and ERβ but with markedly reduced binding affinities and weaker hormonal potency compared to the parent hormone E_2_ (49). However, earlier studies have shown that metabolically-formed 2-OH-E_2_ and 2-OH-E_1_ may be associated with some of the distinct biological functions. For instance, 2-OH-E_1_ was reported to partially antagonize the growth-stimulatory effect of E_2_ in cultured human MCF-7 breast cancer cells (18). This growth-inhibitory effect of 2-hydroxyestrogens (at high concentrations) may be due to their metabolic redox potential to generate reactive estrogen quinones and free radicals which are cytotoxic (18). The enzymatic conversion of 2-OH-E_2_ to 2-methoxyestradiol, which is known to have apoptosis-inducing ability (50–52), may also contribute to its growth-inhibitory effect.

It has been known for many years that the female mice are highly insensitive to APAP-induced liver injury compared to the male mice (44,53). The endogenous estrogens (E_2_ and E_1_) have long been suggested as the confounding factors (53). Some earlier studies have led to the suggestion that an estrogen receptor-dependent mechanism may underlie the hepatoprotective effect of E_2_ and E_1_ against APAP-induced liver injury. The results of our present study show that while E_1_ and E_2_ only have a very weak effect protecting against chemically-induced ferroptosis in cultured hepatoma cells, they exert a similarly-strong protective effect as 2-OH-E_1_ and 2-OH-E_2_ against APAP-induced liver injury in an *in-vivo* mouse model. Given that 2-OH-E_1_ and 2-OH-E_2_are the quantitatively-major metabolites of E_1_ and E_2_ formed in liver by cytochrome P450 enzymes (18), it is reasonable to suggest that the hepatoprotective effect of E_1_ and E_2_ observed *in vivo* likely is mediated, in a significant part, through their metabolic conversion to 2-OH-E_1_ and 2-OH-E_2_, which then exert a strong hepatoprotective effect. In addition, it is known that a small fraction of E_1_ and E_2_ can be metabolically converted to 4-OH-E_1_ and 4-OH-E_2_ (10□15% of 2-OH-E_1_ and 2-OH-E_2_) by hepatic cytochrome P450 enzymes (18). Since 4-OH-E_1_ and 4-OH-E_2_ also have a strong cytoprotective effect against ferroptotic cell death (54), it is thus likely that metabolic formation of 4-OH-E_1_ and 4-OH-E_2_ in the liver may also partially contribute to the observed hepatoprotective effect of E_1_ and E_2_ *in vivo*. Overall, the results of this study offer a novel, estrogen receptor-independent mechanism of cytoprotection against chemically-induced oxidative liver injury both *in vitro* and *in vivo*.

## CONCLUSIONS

As summarized in **Fig. 14**, the results of our present study demonstrate that 2-OH-E_1_ and 2-OH-E_2_, two major endogenous metabolites of E_1_ and E_2_ formed in liver (90), exert a strong protective effect against chemically-induced ferroptosis in liver cells both *in vitro* and *in vivo*, through inhibition of PDI-mediated activation of iNOS/NO, which subsequently results in reduction in cellular ROS and lipid-ROS levels, and ultimately prevention of ferroptotic cell death. In addition, the abrogation of iNOS upregulation by 2-OH-E_1_ and 2-OH-E_2_ and their direct antioxidant activity also partially contribute to their cytoprotective effect in liver cells. These findings reveal a novel mechanism by which 2-OH-E_1_ and 2-OH-E_2_ exert their cytoprotective action in an ER-independent manner. While this study sheds lights on the mechanism underlying the well-known low sensitivity of female rats and mice to APAP-induced hepatotoxicity, it also highlights the therapeutic potential of using endogenous catechol estrogen derivatives for protection against chemically-induced liver injury.

**Figure 14.**
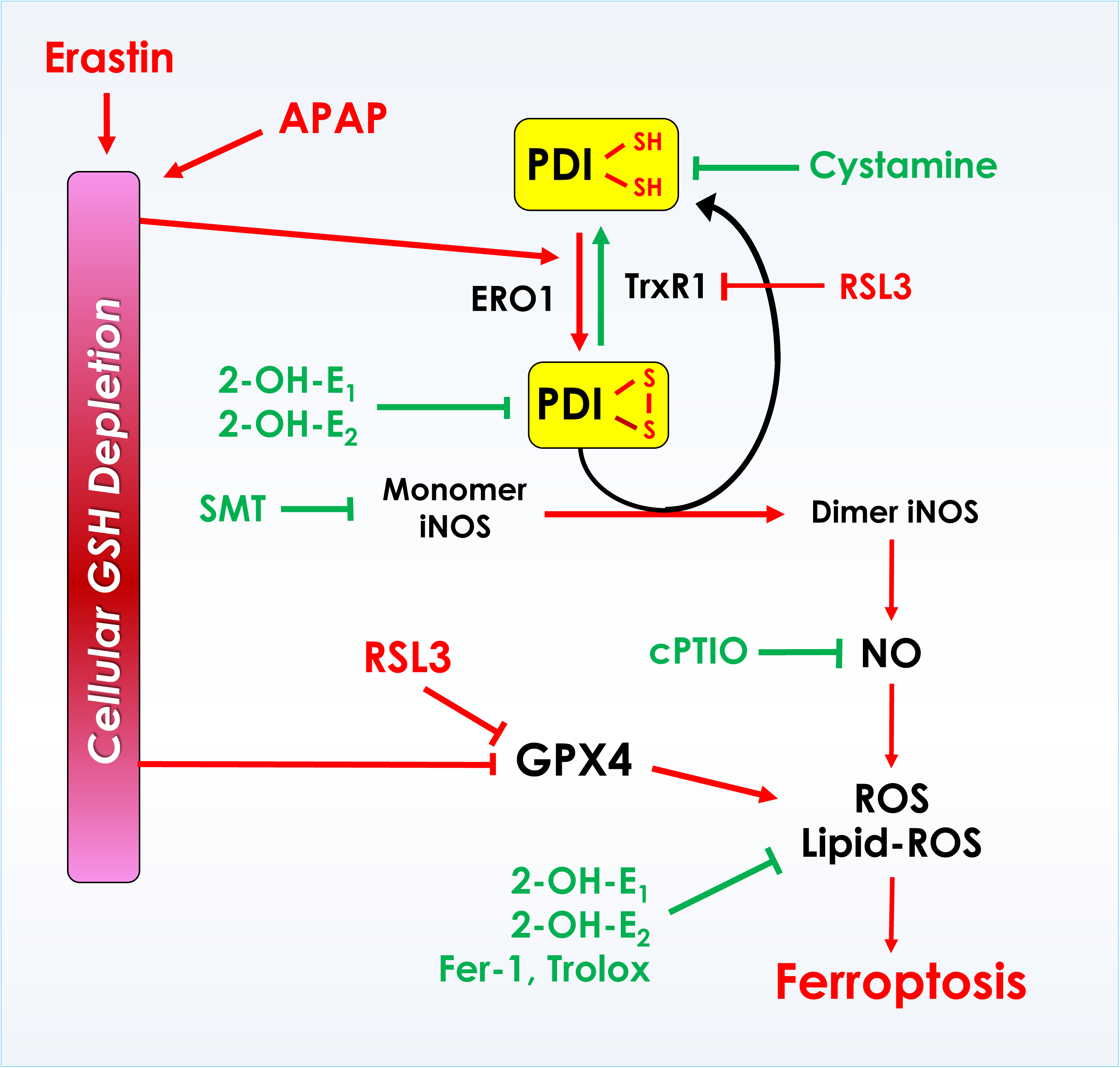
Schematic depiction of the proposed mechanism by which protein disulfide isomerase (PDI) is a target of cytoprotection by 2-OH-E_1_ and 2-OH-E_2_. It is proposed that PDI mediates erastin– and RSL3-induced ferroptotic cell death through PDI-mediated NOS dimerization (activation), which subsequently leads to accumulation of cellular NO and ROS/lipid-ROS and ferroptotic cell death. Inhibition of PDI’s catalytic activity by 2-OH-E_1_ and 2-OH-E_2_ can effectively mediate their protection against chemically-induced ferroptosis in liver cells both *in vitro* and *in vivo*.

## Abbreviations used

E_1_: estrone
E_2_: 17β-estradiol
2-OH-E_1_: 2-hydroxyestrone
2-OH-E_2_: 2-hydroxyestradiol
PDI: protein disulfide isomerase
iNOS: neuronal nitric oxide synthase
iNOS: inducible nitric oxide synthase
NO: nitric oxide
ROS: reactive oxygen species
lipid-ROS: lipid reactive oxygen species

## FIGURE LEGENDS

**Supplementary Figure 1.**
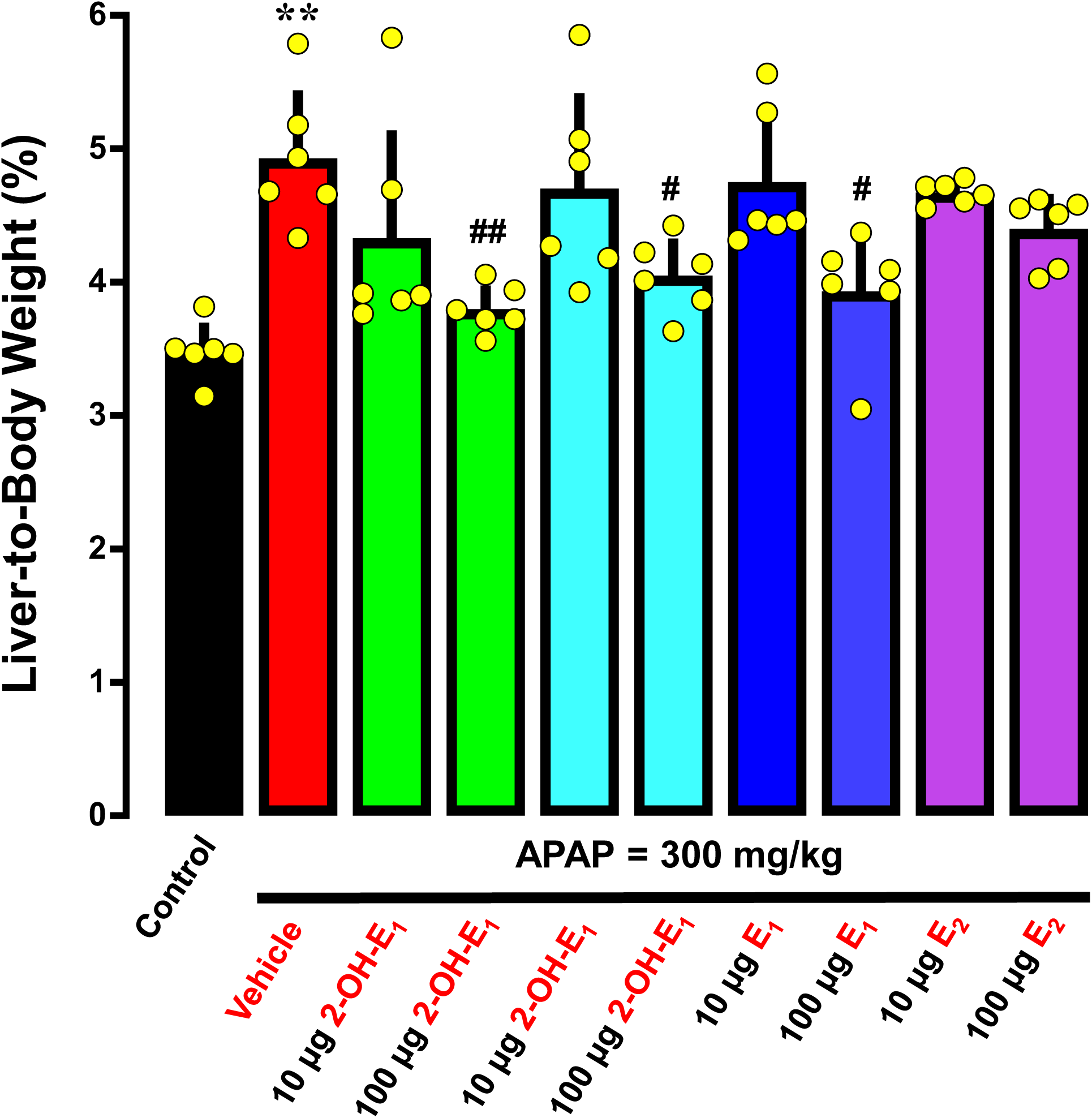
Change in liver-to-body weight ratio in C57BL/6J mice following treatment with APAP ± 2-OH-E_1_, 2-OH-E_2_, E_1_ or E_2_. The animal group and treatment conditions are the same as in Fig. 13 (n = 6). Each value is presented as mean ± S.D. (* or ^#^ *P* < 0.05; ** or ^##^ *P* < 0.01).

